# In vitro programming and pseudounipolarization of human iPSC-derived sensory neurons

**DOI:** 10.64898/2026.09.18.747563

**Authors:** Pascal Röderer, Ana Catarina Costa, Martha Lürkens, Fiona Maria Roll, Franziska Buck, Luzia Heidrich, Jasmin Aicher, Anika Neureiter, Omar Mossad, Holger Trucks-Jansen, Kim Le Cann, Anil Kumar Kalia, Ramya Balaji, Verena Arndt, Andreas Zellner, Yara Mecdad, Thomas Bajaj, Martin Stüger, Martin Karl Schwarz, Anja Nitzsche, Nils Christian Gassen, Natja Haag, Manuel Schröter, Angelika Lampert, Monica M Sousa, Oliver Brüstle

## Abstract

Ectopic expression of *NGN1*, *BRN3A* and *ISLET1* (NBI) from a safe harbor locus in induced pluripotent stem cells (iPSCs) yielded robust differentiation into functional sensory neurons (iNBI-SNs) within seven days. Single nucleus transcriptomics identified peripheral sensory neuron profiles of nociceptors and mechanoreceptors. Electrophysiological studies showed that more than 98 % of iNBI-SNs display TTX-resistant sodium currents and that they establish functional connections to excitatory CNS neurons. iNBI-SNs derived from patients with inherited erythromelalgia, a pain disorder associated with gain-of-function mutations in the Nav1.7 sodium channel showed pathologically increased firing rates which could be partially rescued with a Nav1.7 inhibitor. Notably, iNBI-SNs acquire a characteristic pseudounipolar morphology upon co-culture with embryonic rodent DRG cells. Taken together, NBI-based forward programming of iPSCs represents a robust approach for the generation of human sensory neurons suitable for developmental, disease- and therapy-related studies.

## Introduction

Peripheral sensory neurons are the primary mediators of human somatosensation, and damage to or disease of these neurons is associated with chronic pain and other sensory disorders. They emerge during vertebrate embryonic development from multipotent neural crest cells and can be grouped into three functionally distinct cell types: proprioceptors, mechanoreceptors and nociceptors (Lallemend & Ernfors, 2012; Marmigère & Carroll, 2014). Each of these three main sensory neuron types can be further divided into multiple subtypes, depending on morphological features, projection site within the spinal cord dorsal horn, functional qualities, myelination status and gene expression (Abraira & Ginty, 2013; Chiu et al., 2014; Körner et al., 2026; Li et al., 2016; Tavares-Ferreira et al., 2022; Usoskin et al., 2015; Zheng et al., 2019). Development of these diverse neuronal subtypes is guided via extrinsic signaling, as well as an intricate interaction of a series of transcription factors (Lallemend & Ernfors, 2012; Marmigère & Carroll, 2014; Vermeiren et al., 2020). Neural crest cells develop into sensory neurons in two waves of neurogenesis, initiated by co-expression of Neurog1 (NGN1) and Neurog2 (NGN2), with the first wave mainly being Neurog2- and the second wave being Neurog1-dependent (Ma et al., 1999; Meltzer et al., 2021; Vermeiren et al., 2020). Subsequent co-expression of BRN3A (POU4F1) and ISLET1 stops early neurogenesis and activates a program of terminal differentiation and subtype specification; at the same time expression of non-neuronal genes is repressed (Anderson, 1999; Dykes et al., 2011; Lanier et al., 2009; Sun et al., 2008). While BRN3A and ISLET1 are expressed in all developing sensory neurons, ISLET1 expression was found to be especially important for the development of TrkA and Runx1 expressing nociceptive neurons in mice (Sun et al., 2008). Thus, NGN1/2, BRN3A and ISLET1 constitute a core transcriptional micro-network essential for sensory neuron development. Development of sensory neurons is further associated with a unique morphological transition: Following formation of immature bipolar sensory neurons, they undergo a morphological maturation and change into a pseudounipolar morphology, with a long peripheral and a shorter, thinner central axon branch (Costa et al., 2024; Devor, 1999; Matsuda et al., 2000; Nascimento et al., 2018).

The majority of pain and somatosensation research still relies on animal experiments, despite limited translatability of non-human data to the clinic. The availability of induced pluripotent stem cells (iPSCs) provides a valuable opportunity to generate virtually unlimited numbers of human sensory neurons and thus humanized pain models. This is particularly relevant as human and non-human peripheral neurons have been shown to differ with respect to ion channel expression patterns, specific ion channel functions, as well as subtype specification (Bennett et al., 2019; Bhuiyan et al., 2025; Chang et al., 2018; Jung et al., 2023; Nguyen et al., 2021; Tavares-Ferreira et al., 2022).

Current protocols for the generation of human peripheral sensory neurons mostly rely on the use of small molecules. Chambers et al. reported a strong impact of cell density on *in vitro* differentiation into neural crest cells using dual SMAD inhibition (Chambers et al., 2009; Lee et al., 2010). Although *in vitro* differentiation can be further refined by additional small molecules, factors such as varying cell density remain an issue (Chambers et al., 2012) and can necessitate lengthy adaptation phases for individual iPSC lines (Kalia et al., 2024; Lampert et al., 2020; Röderer et al., 2023). On the other hand, forced overexpression of the transcription factor NGN2 was shown to yield more mature and homogeneous cultures of CNS neurons as compared to small molecule-based approaches (Galiakberova et al., 2025).

Here we use transcription factor-based forward programming for the generation of peripheral sensory neurons. Based on the developmental relevance of *NGN1*, *BRN3A* and *ISLET1* as a differentiation-guiding micro-network, we employ inducible overexpression of these transcription factors to differentiate iPSCs into peripheral sensory neurons. We show that the resulting induced sensory neurons (iNBI-SNs) are electrophysiologically functional, form connections to CNS type neurons, can be employed for disease modeling and become pseudounipolar when co-cultured with DRG glial cells.

## Results

### Overexpression of NGN1, BRN3A and ISLET1 results in efficient differentiation into human sensory neurons

We aimed to identify a transcription factor combination suitable for direct induction of sensory neurons, without transit through an overt intermediate progenitor stage. To this end, we exploited a transcriptional micro-network consisting of NGN1, BRN3A, and ISLET1 (NBI), that governs early *in vivo* patterning of neural crest stem cells towards a sensory neuron fate. In this framework, NGN1 drives acquisition of a neuronal identity, which is subsequently refined by BRN3A and ISLET1 to establish a peripheral sensory neuron phenotype (Fig. 1A). Based on this framework, we hypothesized that the combination of NBI is sufficient to induce differentiation of iPSCs directly into peripheral sensory neurons. Employing genome editing, an inducible transcription factor cassette containing NBI was homozygously integrated into the AAVS1 safe harbor locus of iPSCs (Fig. 1B). Doxycycline-mediated overexpression of these transgenes for seven days induced transition of iPSCs into a neuronal morphology (Fig. 1C-D, Supp. Video 1). Cells were replated one week after initiation of doxycycline treatment in different assay formats and matured in the presence of the neurotrophic factors NGF, BDNF, GDNF and NT-3 to support maturation of sensory neurons (Fig. 1C). In this context we also assessed overexpression of NGN1 alone, as well as a combination of NGN1 and BRN3A. Peripherin (PRPH) expression, a type III intermediate filament protein mainly expressed in neurons of the peripheral nervous system, was used as a first indicator of differentiation success. NGN1 alone generated neuronal cultures with only few PRPH+ neurons, whereas co-expression of NGN1 and BRN3A increased the number of PRPH+ neurons, and additional inclusion of ISLET1 (NBI) further enhanced peripheral sensory neuron differentiation and reduced the amount of non-neuronal cells (Supp. Fig. 1A). Thus, subsequent experiments were focused on iNBI-SN cultures.

**Figure 1.**
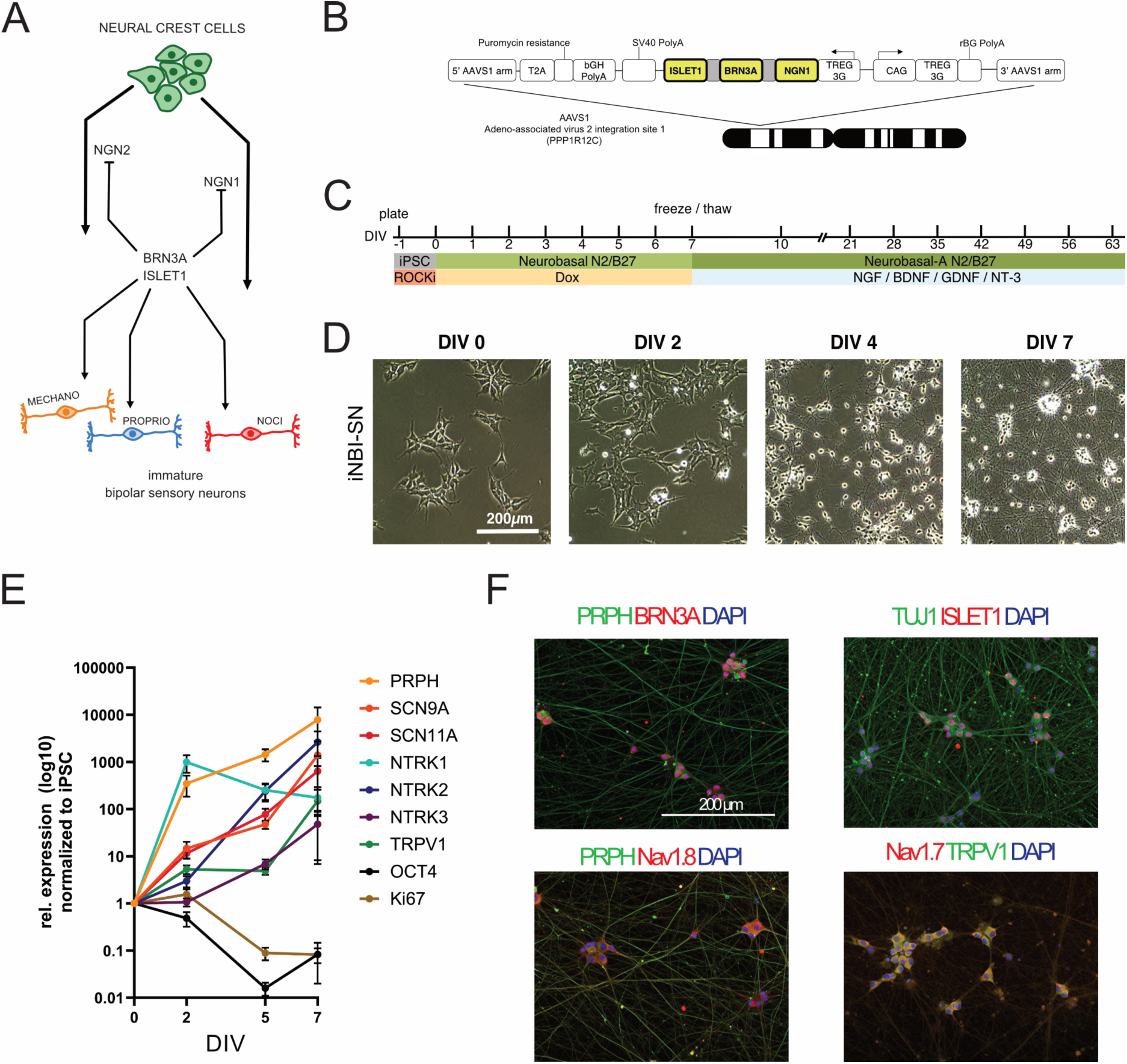
Cellular properties of forward programmed peripheral sensory neurons. **(A)** Simplified representation of the sensory neuron subtype development from multipotent neural crest cells during development. Neural crest cells (NCC) develop in two waves into mechano-, proprio- and nociceptive sensory neurons. Parts of the figure were drawn by using pictures from Servier Medical Art. Servier Medical Art by Servier is licensed under a Creative Commons Attribution 3.0 Unported License (https://creativecommons.org/licenses/by/3.0/). **(B)** Schematic representation of the AAVS1 targeting vector for doxycycline-inducible gene expression. **(C)** Forward programming paradigm for the generation of functional sensory neurons (iNBI-SN). Following transgene induction of *NGN1*, *BRN3A* and *ISLET1* by doxycycline, the cells enter a sensory neuron fate and can be cryopreserved. Further propagation in NGF, BDNF, GDNF and NT-3 is used for full maturation. **(D)** Phase contrast images of the typical morphological changes of iNBI-SN during differentiation. **(E)** Quantitative real-time PCR analysis of gene expression during iNBI-SN differentiation for the sensory neuron marker genes *PRPH*, *SCN9A*, *SCN11A*, *NTRK1, NTRK2, NTRK3* and *TRPV1*, as well as the pluripotency and proliferation marker genes *OCT4* and *Ki67*. Values are presented mean ± SEM, N=12 independent differentiations, pooled from two healthy donors **(F)** Immunocytochemical staining of sensory neuron markers BRN3A, ISLET1, TUJ1, Peripherin, TRPV1, Nav1.7 and Nav1.8 in iNBI-SN cultures at DIV 63.

During overexpression of NBI we observed a consecutive downregulation of *Ki67* and *OCT4*, marker transcripts associated with proliferation and pluripotency, respectively. Conversely, expression of marker genes of a peripheral sensory neuron identity such as *PRPH*, *SCN9A*, SCN11A, *NTRK1*, *NTRK2*, *NTRK3* and *TRPV1* gradually increased from DIV 2 onwards (Fig. 1E). Expression of *PRPH* and *NTRK1* had already increased about 350- and 990-fold, respectively by DIV 2 indicating a very early fate entry (Fig. 1E). Immunofluorescence analyses revealed that expression of BRN3A, ISLET1, PRPH, Nav1.7, Nav1.8 and TRPV1 was retained until at least eight weeks after replating (Fig. 1F, Supp. Fig. 1B).

To obtain further insight into the dynamics of fate transition, we performed bulk RNA sequencing across the timeframe of transcription factor overexpression. Principal component analysis confirmed that iNBI-SNs cluster together according to the time point of harvest with DIV 7 clustering separately from DIV 35 and 63, along PC1 (Fig. 2A), substantiating the notion that iNBI-SNs mature over time with an initial rapid maturation, that then slows down. Along this line, differential gene expression analysis showed that the number of DEGs increases markedly from DIV 7 to DIV 35 and DIV 65, whereas only a comparatively small number of DEGs was detected between DIV 35 and DIV 63 (Supp. Fig. 2A-C). Clustering of differentially expressed genes identified three main gene clusters in the differentiating iNBI-SN cultures over time (Fig. 2B). To functionally characterize these clusters, a gene ontology (GO) enrichment analysis was performed using biological process terms, and the top 10 GO terms, ranked by GeneRatio were visualized (Fig. 2C). Cluster 1, comprising genes that show a high expression at DIV 7 and are then downregulated at DIV 35 and 63, was significantly enriched for GO terms associated with neuronal development, such as axogenesis, synapse organization, sensory system development and action potential, suggesting an early immature developing sensory neuron phenotype at DIV 7. Clusters 2 and 3, representing genes upregulated at DIV 35 and DIV63, respectively, were enriched for GO terms associated with neuronal function such as modulation of chemical synaptic transmission, regionalization, positive regulation of MAPK cascade, suggesting a functional refinement over time. In addition, GO terms indicating an ongoing development such as regulation of nervous system development and sensory system development were also identified in these clusters.

**Figure 2.**
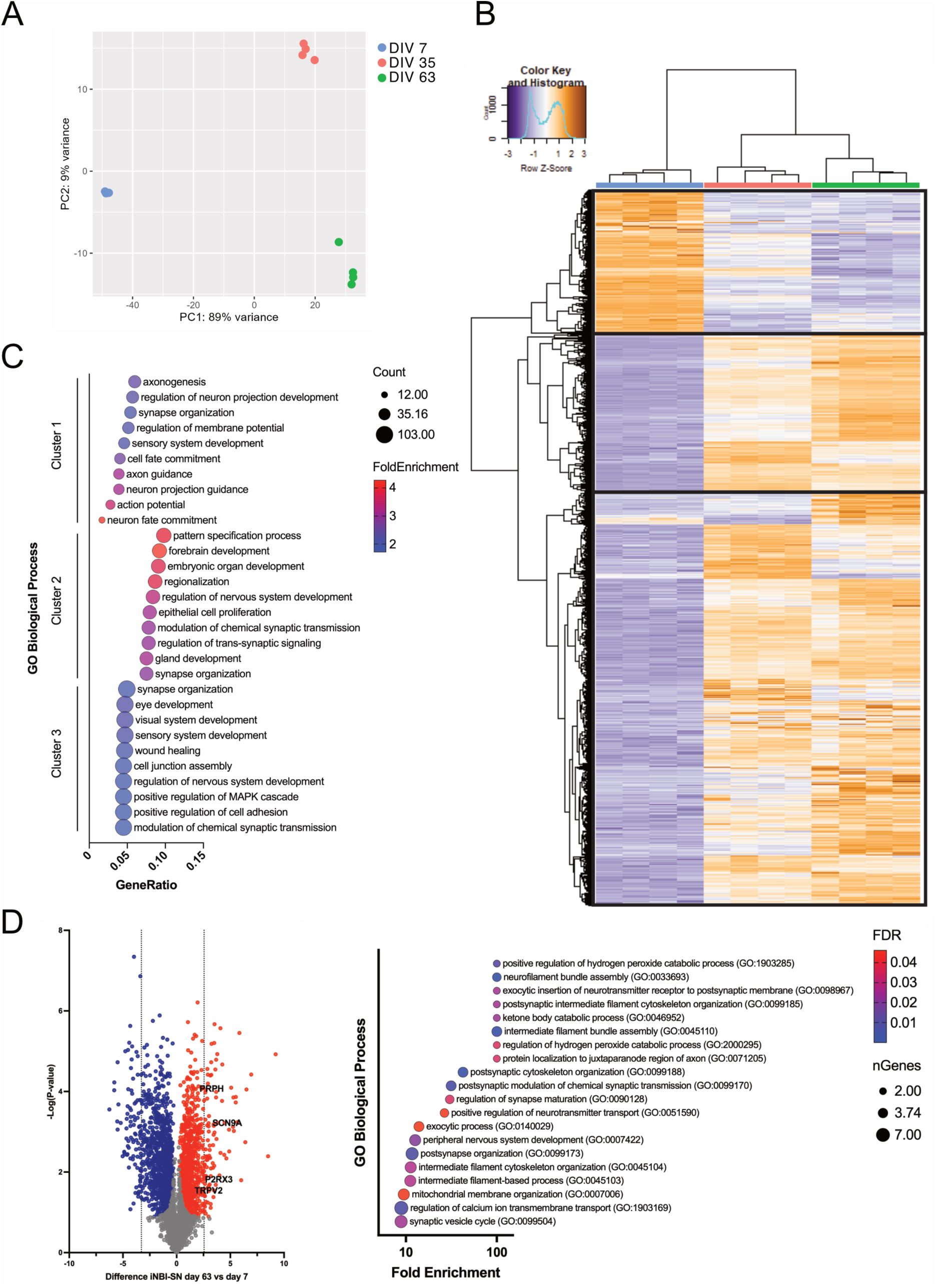
Dynamic gene and protein expression changes in iNBI-SNs. **(A)** Principal component analysis (PCA), illustrating the clustering of bulkRNAseq data across two principal components, accounting for 89.0 % (PC1) and 9.0 % (PC2) of total variance, indicating that time is the main driver of variance. PCA was performed, employing FPKM normalized gene expression data. **(B)** Heatmap showing the expression profile of differentially expressed genes across the time course of in vitro differentiation (DIV7, 35 and 63). Boxes mark three identified gene clusters. **(C)** Gene Ontology enrichment analysis for biological processes, based on the three gene clusters identified in the heatmap. **(D)** Volcano plot comparing protein expression levels on DIV 7 and 63 of differentiation. Significantly upregulated proteins are highlighted in red, significantly downregulated proteins are labeled in blue. Dotted lines mark the top 100 up- and downregulated proteins. Amongst the upregulated proteins are Peripherin (PRPH), Nav1.7 (SCN9A), P2RX3 and TRPV2 indicating a structural and functional maturation of sensory neurons over time in culture. Gene Ontology enrichment analysis for biological processes, based on the top 100 upregulated proteins on DIV 63 in iNBI-SNs.

Analyzing protein expression at DIV 7 and DIV 63 corroborated these findings. We performed a GO overrepresentation analysis of the 100 most upregulated proteins in iNBI-SN at DIV 63 compared to DIV 7 to identify biological processes significantly enriched within our dataset. Similar GO terms as previously identified in the transcriptome data, such as peripheral nervous system development, regulation of synapse maturation and neurofilament bundle assembly were identified underlining the functional and structural maturation of iNBI-SNs over the time of in vitro culture (Fig. 2D). In this context we also compared the proteome signature of iNBI-SN to sensory neurons generated via overexpression of NB and sensory neurons, that were differentiated via a classical small molecule protocol (Suppl. Fig. 2D). Principal component analysis of protein expression revealed a clear separation between conditions and time points (Suppl. Fig. 2D). At the early time point (immature neurons), classically differentiated sensory neurons formed a distinct and distant cluster, whereas both forward programmed sensory neuron populations clustered closer together and interspersed between early classically differentiated SNs and all samples of late DIV stages (matured neurons). These results suggest that forward programming of sensory neurons promotes a more mature proteomic state already at early stages. In contrast, at the late time points (DIV 63 and DIV 70) all cell types converged towards a single cluster, indicating an increased similarity in protein expression profiles after prolonged culture (Suppl. Fig. 2D).

Together this data establishes NBI forward programming as a suitable strategy for the rapid generation of DRG-like sensory neurons from iPSCs, yielding cells that exhibit an accelerated maturation versus conventional differentiation protocols while maintaining a comparable expression profile.

### Single cell analysis reveals enrichment of mechano- and nociceptive sensory neurons in iNBI-SN cultures

Bulk transcriptomic analysis indicated that NBI-induced cells co-express multiple sets of peripheral sensory neuron-specific ion channels and receptors. To determine whether this reflects a homogeneous expression profile or the presence of distinct cellular subpopulations, we performed RNAscope using a panel of probes targeting sensory neuron-specific transcripts (Fig. 3A). Co-expression patterns were analyzed across five panels, each comprising three marker genes representing ion channels, receptors and neuropeptides characteristic of distinct sensory neuron subtypes (Fig. 3A). In order to assess developmental changes this analysis was performed at DIV 35 and DIV 63. Probing expression of the archetypical voltage gated sodium channel genes *SCN9A*, *SCN10A* and *SCN11A* at DIV 35, the largest fraction of iNBI-SNs expressed *SCN9A* (34.14 %), while 19.35 % of cells co-expressed all three ion channels. By DIV 63, co-expression of all three channels increased markedly to 42.20 %, indicating a progressive maturation of nociceptor-like profiles. A similar increase in co-expression over time was observed for the combinations *PIEZO2*, *CALCB* and *TRPM8* (markers associated with low-threshold mechanoreceptors (LTMRs) and nociceptors), as well as *PENK*, *OSMR* and *SST* (markers associated with itch nociceptors, PENK+ nociceptors and sleeping nociceptors). In contrast, the combinations *ASIC1*, *NTRK2* and *NTRK3* (associated with LTMRs and proprioceptors) and *TAC1*, *HTR3A* and *NTRK1* (associated with peptidergic nociceptors) showed more segregated expression patterns, with a reduced co-expression at the later time point. Beyond changes in co-expression, gene expression dynamics also varied at the single-gene level. Four distinct patterns emerged: (i) increased mean expression levels accompanied by an increased proportion of expressing cells (*SCN10A, SCN11A, CALCB, SST, OSMR*), (ii) increased mean expression levels without a change in number of expressing cells (*HTR3A, TAC1, PIEZO2, PENK*), (iii) increased mean expression levels with a reduced proportion of expressing cells (*NTRK3, TRPM8*), and (iv) decreases both in mean expression levels and number of expressing cells (*SCN9A, ASIC1, NTRK1, NTRK2*) (Suppl. Fig. 3). Overall, this data indicates a dynamic and time-dependent refinement of gene expression, consistent with progressive diversification into transcriptionally more distinct sensory neuron subtypes during *in vitro* maturation.

**Figure 3.**
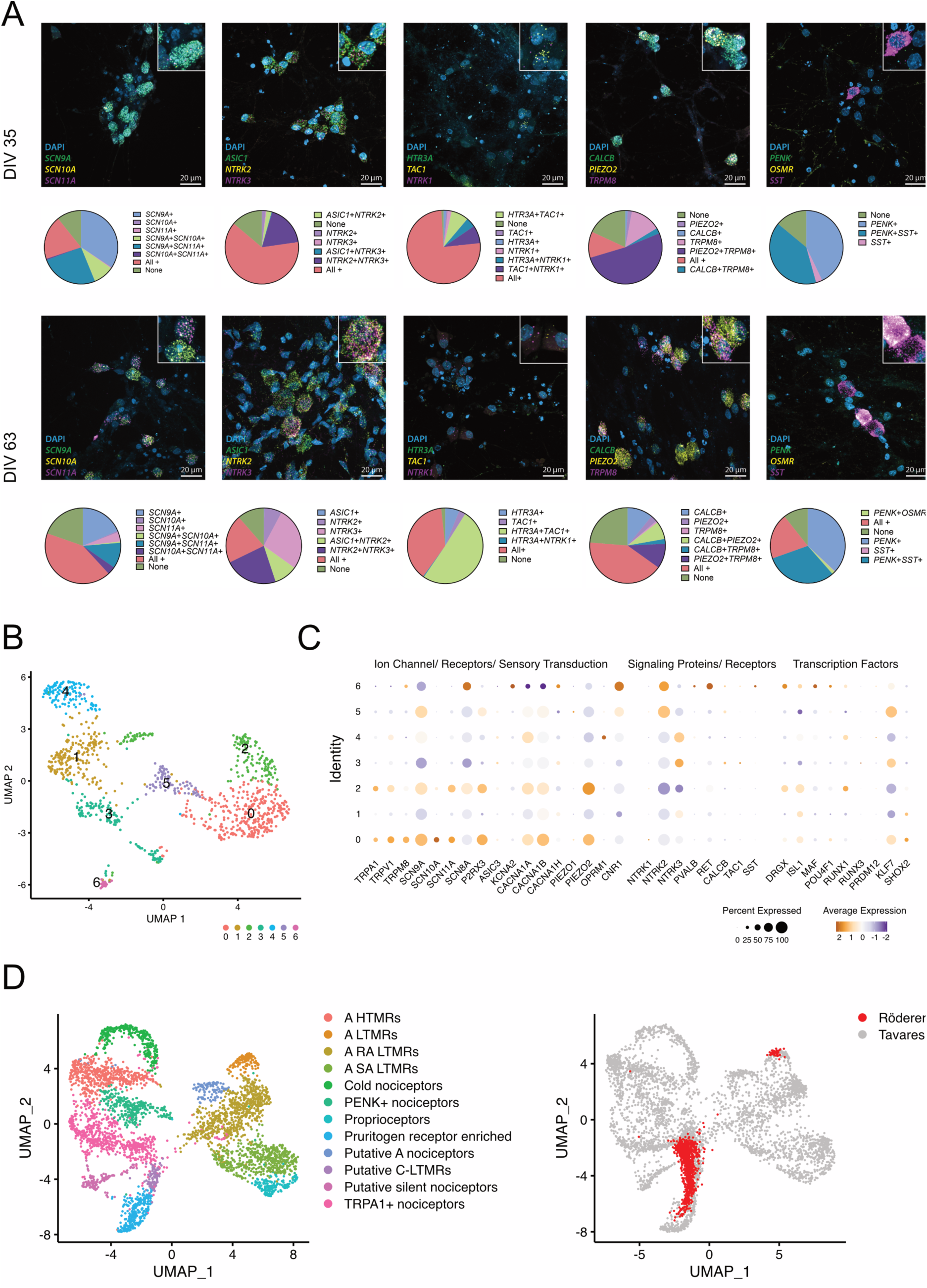
Sensory neuron subtype specification in iNBI-SNs. **(A)** RNAscope in situ hybridization analysis. Merged images for the combination of each three targets and DAPI at DIV 35 and 63 are shown. Inserts show positive cells in higher magnification. Population distribution of marker combinations is shown in pie charts below. **(B)** UMAP subclustering of mature neurons extracted from the whole population UMAP (Suppl. Fig. 4A) at DIV 63 based on two genetic backgrounds. **(C)** Dot plot showing the expression of genes and markers representing ion channels, signaling proteins and transcription factors across clusters. **(D)** Comparison of transcriptomic signature of iNBI-SN and adult human DRG neurons. Left: UMAP plot showing the 12 human DRG neuronal clusters identified by Tavares-Ferreira et al., 2022 (Tavares-Ferreira et al., 2022). Right: UMAP with iNBI-SN nuclei (red dots) projected onto the adult DRG Atlas based on transcriptomic similarity, indicating the presence of A-LTMRs, pruritogen receptor enriched, putative C-LTMRs and TRPA1+ nociceptors.

Segregation in distinct neuronal subpopulations was further confirmed by single nucleus RNA sequencing. Unsupervised clustering of the single-nucleus dataset identified 10 distinct cell clusters. GO analysis of cellular component terms revealed enrichment for diverse biological features, including cell adhesion, extracellular matrix, mitochondrial, ribosomal and synaptic structures (Supp. Fig. 4A-B). Neuronal markers were detected across all clusters, but were most prominently expressed in clusters enriched for neuronal and synaptic GO terms, including asymmetric synapse, neuron-to-neuron synapse and synaptic membrane. To focus on the most mature neuronal population, nuclei of these clusters were subclustered into seven distinct clusters (Fig. 3B). Marker gene analysis indicates that clusters 0 and 2 were enriched for canonical nociceptor-associated transcripts, including *TRPV1, TRPA1*, as well as the voltage-gated sodium channels *SCN10A* and *SCN11A*. Clusters 5 and 6 were enriched for *NTRK2* expression, while clusters 3 and 4 showed stronger expression of *NTRK3*, markers associated with a mechano- and proprioceptive identity, respectively (Fig. 3C). Assignment to defined sensory neuron subtypes remained challenging due to the extensive overlap and co-expression of subtype markers, consistent with our RNAscope findings. Therefore, the subtype identities of these *in vitro*-generated neurons were inferred by comparing their transcriptional signatures to a spatial transcriptomics atlas of the *in vivo* adult dorsal root ganglion (Tavares-Ferreira et al., 2022). This mapping indicates highest transcriptional similarity of iNBI-SNs with A-LTMRs and TRPA1+ nociceptors, and to a lesser extend to putative C-LTMRs and pruritogen receptor enriched neurons (Fig. 3D, Supp. Fig. 4C). Collectively, these data demonstrate that NBI-driven differentiation gives rise to a mixed culture of relevant sensory neurons, enriched for mechano- and nociceptive subtypes.

### ‘NBI’ derived sensory neurons show functional properties of peripheral nociceptive neurons

#### iNBI-SNs express functional TTX-resistant sodium channels

We next analyzed iNBI-SNs by whole-cell current-clamp recordings at DIV 21, 35, 49 and 63, using cells from two independent genetic backgrounds, with both cell lines showing consistent results (Fig. 4). The majority of analyzed neurons showed a tonic firing pattern upon current injection (Fig. 4A-B), and only 7.5 – 15 % of neurons exhibited spontaneous action potential discharges at very low frequencies, as is typically observed in C-fibers of healthy individuals (Fig. 4C). The resting membrane potential of iNBI-SNs gradually hyperpolarized from −62.31 ± 0.99 mV at DIV 21 to −72.46 ± 1.14 mV (mean ± SEM, Fig. 4D). The rheobase and action potential peak amplitude gradually increased over time from 40.88 ± 3.23 pA to 122.10 ± 8.57 pA and from 88.30 ± 1.65 mV to 111.80 ± 2.26 mV, respectively (mean ± SEM, Fig. 4D). The action potential half-width decreased from 3.17 ± 0.14 ms at DIV 21 to 2.78 ± 0.16 ms at DIV 49, to then increase again to 3.36 ± 0.22 seconds at DIV 63 (mean ± SEM, Fig. 4D). Similarly, the action potential firing behavior evolved over time. At earlier stages (DIV 21 and 35), iNBI-SNs showed a steep increase in action potential firing frequency in response to increasing current injections, which declined at higher current amplitudes, potentially indicating an earlier onset of a depolarization block. In contrast, at later stages (DIV 49 and 63), iNBI-SNs exhibited a more gradual increase in firing frequency with increasing current, which eventually plateaued (Fig. 4E). We further analyzed sodium currents, as peripherally expressed sodium channels Nav1.7, Nav1.8 and Nav1.9 are not only employed as subtype markers, but are also of special interest in pain pharmacology. Specifically, we investigated tetrodotoxin-sensitive (TTXs) and -resistant (TTXr) sodium currents. Interestingly, the majority of the sodium current was found to be mediated by TTXs channels; more than 98 % of analyzed neurons showed TTXr current components across both cell genetic backgrounds (Fig. 4F). Application of 3 µM VX-150, a specific Nav1.8 inhibitor, allowed the partial attribution of the TTXr currents to the peripheral TTXr sodium channel Nav1.8 (Suppl. Fig. 5A), while no Nav1.9-like persistent current could be identified. Overall, the sodium current profile was comparable between individual cells, with slight variation in current proportions (Suppl. Fig. 5B). Additionally, application of a specific Nav1.7 inhibitor SSCI-2, abolished spontaneous firing in iNBI-SN cultures (Suppl. Fig. 5C). Taken together, these data demonstrate a progressive functional maturation of iNBI-SNs over time in culture, including the recapitulation of key features of peripheral sensory neurons.

**Figure 4.**
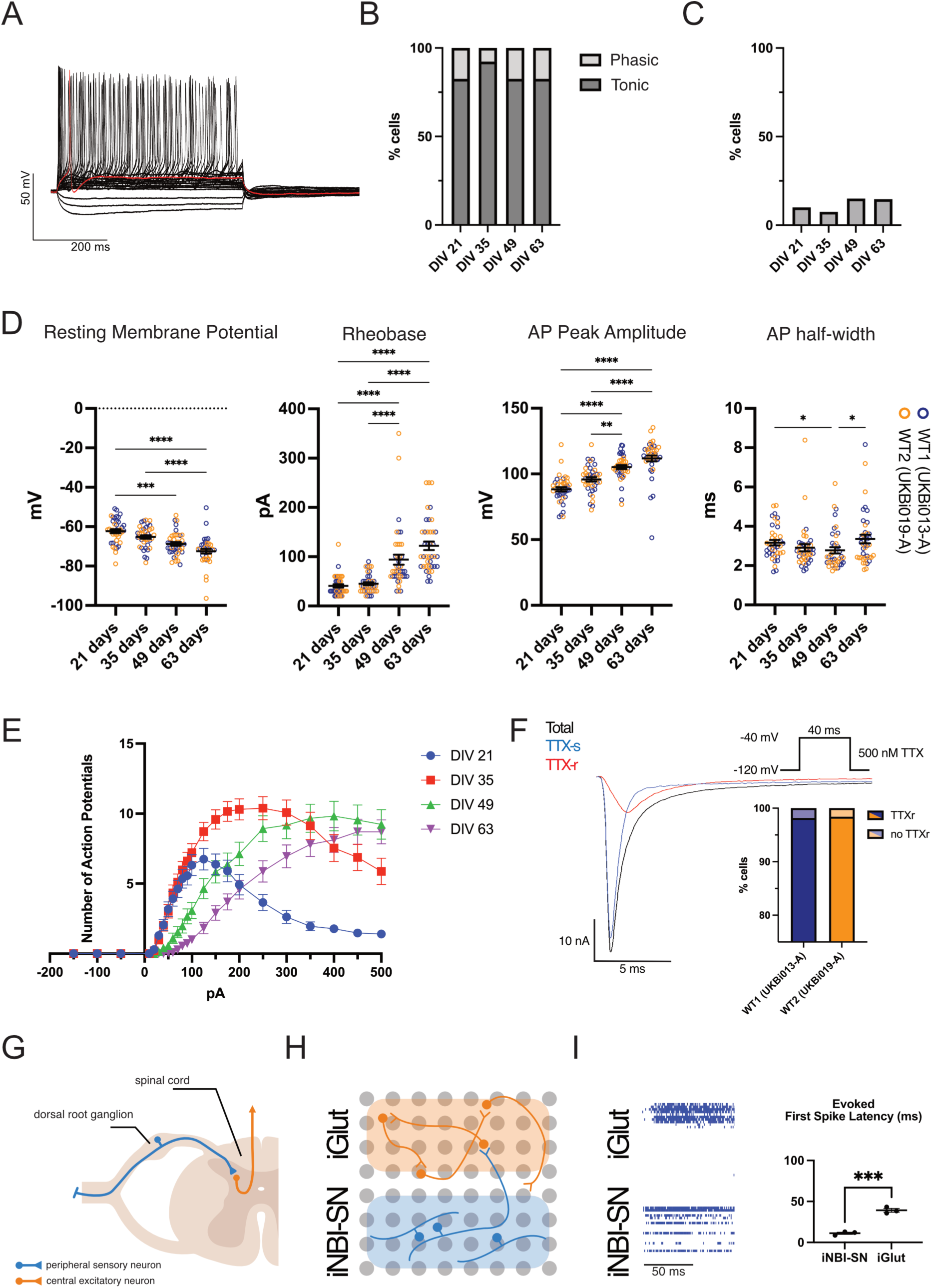
Development of electrophysiological properties of iNBI-SNs over time. **(A)** Exemplary trace of induced action potential (AP) firing in iNBI-SN at DIV 63, red trace indicates AP at rheobase. **(B)** Quantification of phasic and tonic firing behavior of iNBI-SNs over time. N=39-40 cells per time point, pooled from the two wildtype cell lines UKBi013-A and UKBi019-A. **(C)** Quantification of spontaneously active neurons in iNBI-SN cultures over time. N=39-40 cells per time point, pooled from the two wildtype cell lines UKBi013-A and UKBi019-A. **(D)** Basic electrophysiological parameters (resting membrane potential, rheobase, AP peak amplitude, AP half-width) develop over the time of in vitro culture. N=39-40 cells per time point, pooled from the two wildtype cell lines UKBi013-A and UKBi019-A. Values are presented as mean ± SEM with single data points indicated by circles. Kruskal-Wallis test with Dunn’s multiple comparisons test. **(E)** Action potential firing behavior in response to a series of 500 millisecond current steps from −150 to 500 pA. Data are presented as mean ± SEM. **(F)** Exemplary sodium current trace before and after application of 500 nM TTX, indicating the TTX-resistant (red) and sensitive (blue) sodium current. More than 98 % of iNBI-SNs showed TTX-resistant currents at DIV 63. N=55 cells from UKBi013-A and N= 63 from UKBi019-A. **(G)** Schematic of peripheral sensory neuron interaction with the neural circuitry in the spinal cord dorsal horn. **(H)** Schematic overview of iNBI-SN and iGlut multi-electrode connectivity assay. **(I)** Raster plot of evoked spiking behavior upon optical stimulation. Optogenetic activation of ChRmine elicits electrical activity in iNBI-SNs, that is then transmitted to iGlut neurons. Quantification of the evoked first spike latency upon optical stimulation. N=3 wells with 32 electrodes / neuron type / well. Values are presented as mean ± SEM. Unpaired t-test.

#### iNBI-SNs establish functional connections to CNS neurons

One of the main tasks of peripheral sensory neurons is to collect information from their target tissues and forward this information towards the brain via the spinal cord dorsal horn (Fig. 4G). To validate iNBI-SNs in this regard we used side-by-side cultures with CNS neurons on multi-electro-arrays (MEAs). To that end we first transduced our iNBI-SNs with an adeno-associated virus (AAV) carrying the optogenetic activator channel ChRmine coupled to a mScarlet fluorescent reporter (Suppl. Fig. 6). As recipient population we employed NGN2-induced glutamatergic (iGlut) neurons as a proxy for the CNS compartment (Peitz et al., 2020; Rhee et al., 2019; Y. Zhang et al., 2013). Both populations were plated spatially separated next to each other on a MEA chip (Fig. 4H). Upon optical stimulation of iNBI-SNs, light stimulation-induced activity could indeed be seen propagating to the adjacent iGlut neurons (Fig. 4I). While the iNBI-SN compartment showed an evoked response probability of close to 99%, the iGlut neuron compartment still showed a response probability of 89% (Suppl. Fig. 6D). The number of evoked spikes did not significantly differ between both compartments, while the iNBI-SN compartment showed a significantly lower latency of the first spike, compared to the iGlut neurons (Fig. 4I, Suppl. Fig. 6D). In absence of ChRmine in the iNBI-SN compartment only random signals were recorded (Suppl. Fig. 6D). These data demonstrate the iNBI-SNs can establish functional connectivity to CNS neurons and engage in directional signal propagation.

#### iNBI-SNs are amenable to disease modeling and compound-mediated rescue

To explore the amenability of iNBI-SN cultures to disease modeling and drug testing we generated neurons from two independent healthy donors, as well as two independent genetic chronic pain patients. The patients were diagnosed with inherited erythromelalgia (IEM), which is associated with severe attacks of burning pain, redness and swelling of the distal extremities and caused by mutations in the SCN9A gene coding for the peripheral sodium ion channel Nav1.7. The two patients carried distinct gain-of-function mutations (IEM1: V400M, IEM2: F1449V). IEM patient derived iNBI-SNs showed an increased firing behavior, which became more pronounced over time (Fig. 5A-B). At DIV 63 the mean firing rate of both control cell lines was 1.18 ± 0.49 Hz and 1.69 ± 0.46 Hz, while both patient specific cell lines showed increased firing rates of 4.82 ± 0.46 Hz and 3.88 ± 0.71 Hz (Fig 5C). The mean firing rate clearly distinguished wildtype and patient derived cells. IEM1 showed a significantly increased number of bursts, compared to WT1 while IEM2 only showed a trend to an increased bursting behavior (Fig. 5C). Interestingly, the bursts themselves (i.e. burst duration, number of spikes per burst, inter-burst-interval) did not differ between WT and IEM lines (Fig. 5C). Although currently no selective Nav1.7 inhibitor is in clinical use, multiple inhibitors were developed and preclinically tested. Here we applied the Nav1.7 inhibitor PF-05089771 at 200 nM and 1000 nM for its potential to decrease the pathological hyperactivity in IEM patient derived iNBI-SNs. Application of PF-05089771 significantly reduced the mean firing rate, as well as the number of bursts in patient specific iNBI-SNs (Fig. 5D-E). Together, these findings demonstrate that iNBI-SNs can capture disease-associated changes in neuronal activity and can be effectively used for pharmacological testing of drugs and other compounds.

**Figure 5.**
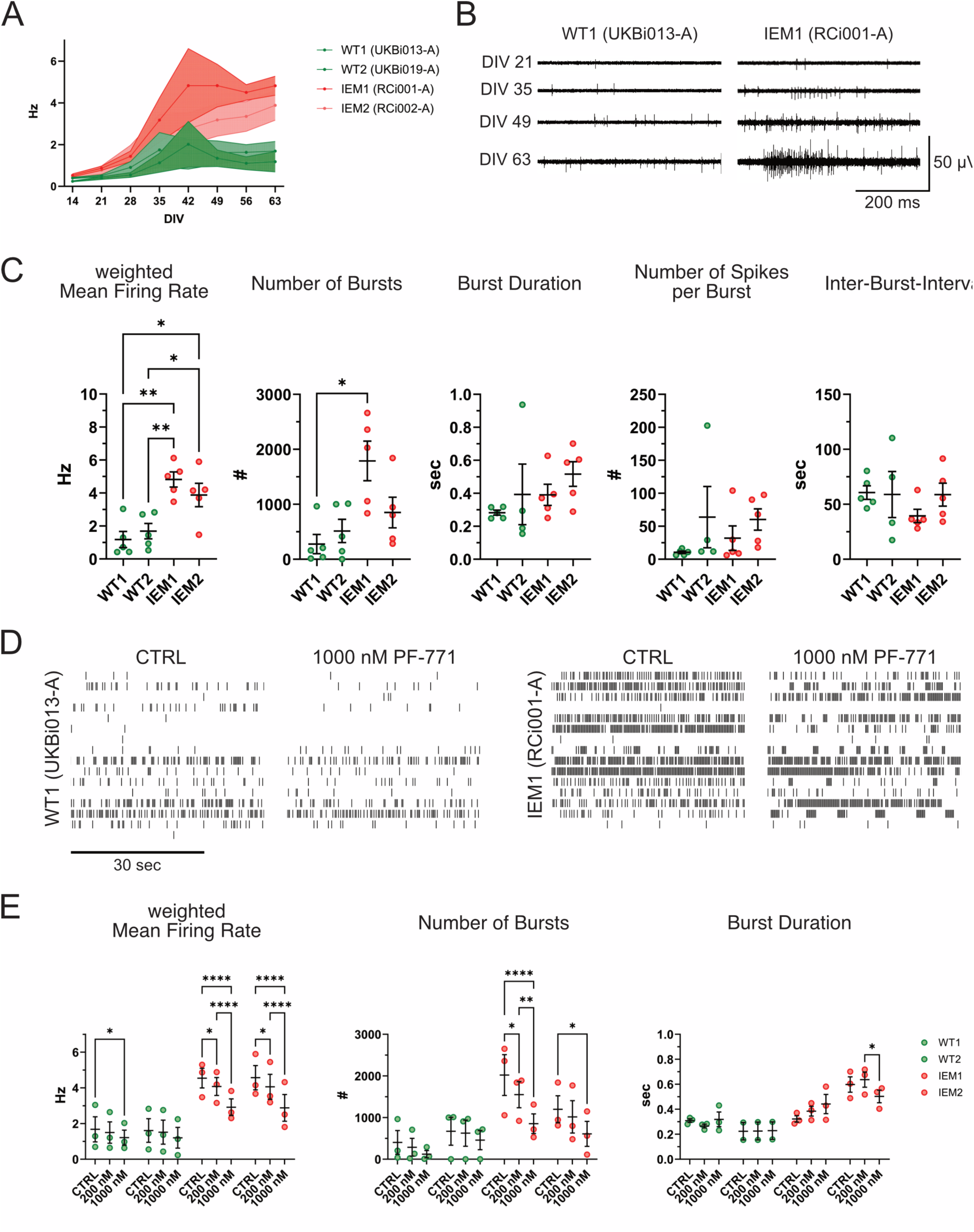
Modelling genetic pain disorder: inherited erythromelalgia. **(A)** Development of spontaneous firing behavior and disease phenotype over time of in vitro culture. Quantification of weighted mean firing rate during 15 minutes multi-electrode array recordings performed weekly between DIV 14 and 63. Data is presented as mean ± SEM. N=5 independent experiments, with 6 wells per experiment. **(B)** Exemplary voltage traces of wildtype (UKBi013-A) and patient (RCi001-A) derived iNBI-SNs at DIV 21, 35, 49 and 63. **(C)** Quantification of weighted mean firing rate, number of bursts, burst duration, number of spikes per burst and inter-burst-interval during 15 minutes multi-electrode array recordings at DIV 63. N=5 independent experiments, with 6 wells per experiment. Values are presented as mean ± SEM with single data points indicated by circles. Kruskal-Wallis test with Dunn’s multiple comparisons test. **(D)** Representative raster plots of spontaneous firing behavior of WT1 and IEM1 iNBI-SN cultures at DIV 63 before (CTRL) and after application of 1000 nM PF-05089771 (PF-771), a selective Nav1.7 inhibitor. **(E)** Quantification of weighted mean firing rate, number of bursts and burst duration during 15 minutes multi-electrode array recordings at DIV 63 at baseline and after application of 200 nM and 1000 nM PF-05089771. N=3 independent experiments, with 5-6 wells per experiment. Values are presented as mean ± SEM with single data points indicated by circles. Repeated measurement two-way ANOVA with Tukey’s multiple comparisons test.

### Co-culture of iNBI-SNs with rat DRG glial cells induces an authentic pseudounipolar morphology with asymmetric structure and function

Acquisition of a pseudounipolar morphology with a central and a peripheral axon branch emanating from a joint stem axon is a hallmark of human peripheral sensory neurons. Commonly employed protocols for the generation of human sensory neurons from iPSCs lack a mature pseudounipolar morphology. Recent rodent data show that the morphological maturation from a bipolar into a pseudounipolar morphology can be triggered by co-culture and cell-cell interaction with dorsal root ganglion glial cells (Nascimento et al., 2022). When we co-cultured iNBI-SNs from DIV 7 onwards with E16 rat DRG glial cells (Fig. 6A), we observed emerging pseudounipolarized neurons as well as bipolar and bellshaped-bipolar neurons transitioning towards pseudounipolarity (Fig. 6B). Live imaging of the morphological transition from a multipolar neuron into a pseudounipolar morphology indicates that initially supernumerary axons are retracted, while the presumptive central and peripheral axons move towards each other to then establish a joint stem axon (Fig. 6C, Suppl Video 2). This morphological maturation was observed to be a relatively fast process occurring over a time period of about 12 hours and was reversible in some neurons. To test if the morphological maturation is triggered by factors released from peripheral glial cells or via direct cell-cell interaction we cultured iNBI-SNs under control conditions, with filtered glial-conditioned medium and in direct co-culture. While under control and filtered medium conditions no morphological maturation was observed, direct co-culture showed induction of a pseudounipolar morphology in 25.77 ± 4.70 % of neurons (Fig. 6D), indicating the need for a direct cell-cell interaction. Upon closer inspection, we found a morphological asymmetry in pseudounipolarized neurons, always presenting with a thicker and a thinner axon, resembling a peripheral and a central axon branch respectively (Fig. 6E).

**Figure 6.**
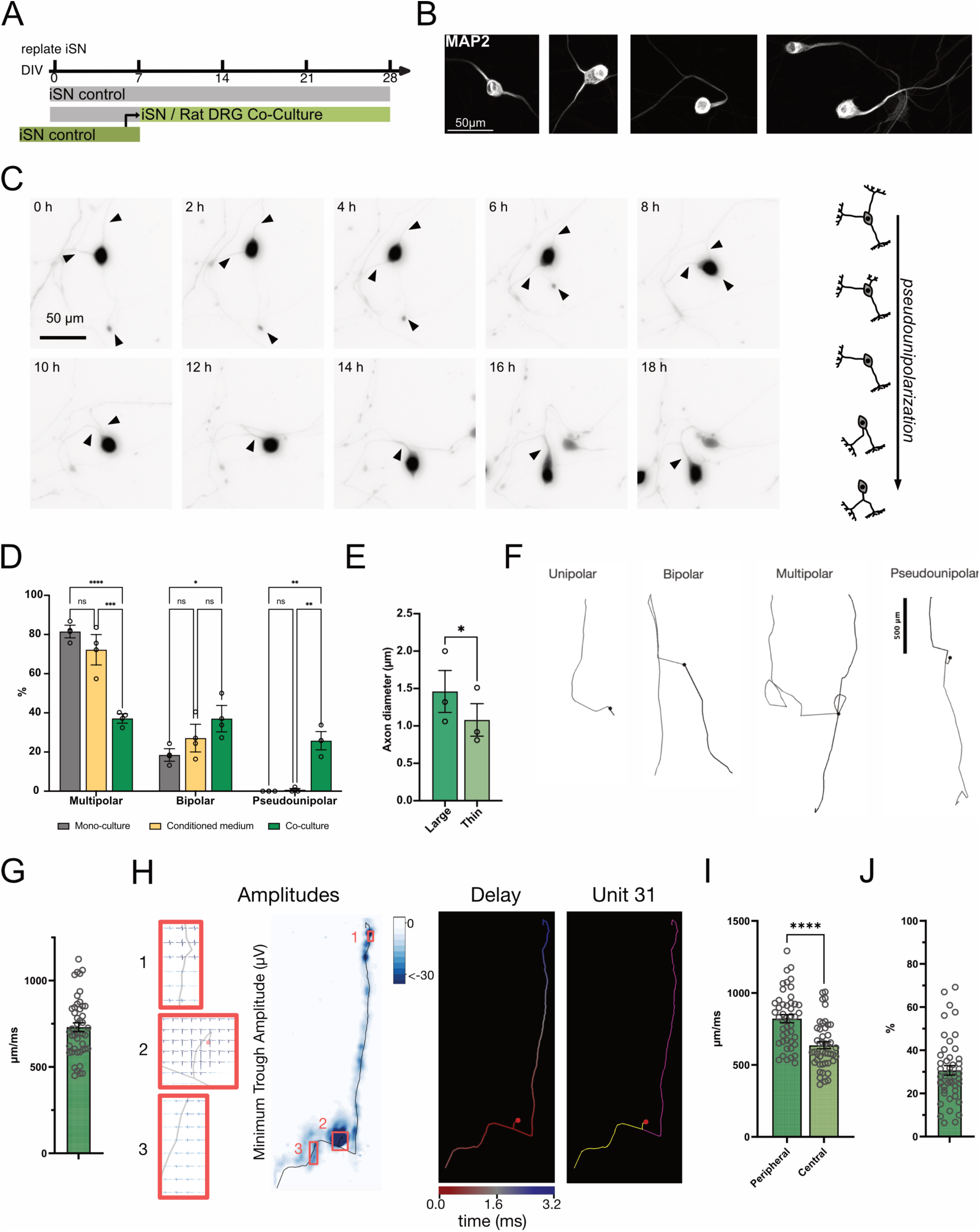
Co-culture of iNBI-SNs with rat dorsal root ganglia cells induces a pseudounipolar morphology. **(A)** Co-culture paradigm for induction of pseudounipolarization in iNBI-SN. **(B)** Representative images of iNBI-SN at different stages of pseudounipolarization after DIV 28. **(C)** Live-imaging of tdTomato-expressing iNBI-SNs during the process of pseudounipolarization in co-culture with rat DRG cells. Arrows indicate individual axons throughout this process. A supernumerary axon present at 0 h progressively retracts and is absent by 8 h. The two remaining axons subsequently move towards each other and merge to form a single pseudounipolar stem axon by 12 h. Adjacent schematic illustrates the observed morphological transition. **(D)** Quantification of sensory neuron morphologies after three weeks of mono-culture (control), treatment with conditioned medium or direct co-culture. N=3 experiments, with 70 to 180 cells per experiment. Values are presented as mean ± SEM with single data points indicated by circles. Two-way ANOVA with Tukey’s multiple comparison test. **(E)** Axon diameter of pseudounipolar iNBI-SNs displaying a large and thin axon. N= 3 independent experiments. Paired t-test. **(F)** Reconstructions of the axonal morphology from electrical footprints of iNBI-SNs in co-culture with rat DRG cells on the HD-MEA. Shown are four morphology types identified in the cultures. **(G)** Average conduction velocity of pseudounipolar iNBI-SNs. N=46 units pooled across 6 wells, 2 independent experiments. Values are presented as mean ± SEM with single data points indicated by circles. **(H)** Exemplary axon scan of a pseudounipolar iNBI-SN during a HD-MEA recording indicating amplitudes, signal delay and morphology. **(I)** Quantification of peripheral (larger) and central (thiner) axonal branch conduction velocities in pseudounipolar iNBI-SNs. N=46 cells. Values are presented as mean ± SEM with single data points indicated by circles. Paired t-test. **(J)** Quantification of maximum velocity difference (%) between peripheral and central axons. N=46 cells. Values are presented as mean ± SEM with single data points indicated by circles.

Next, we applied high-density microelectrode array (HD-MEA) recordings to investigate the detailed morpho-electrical features of pseudounipolarized human sensory neurons. Following axon tracking across the whole HD-MEA, spike-sorting, and spike-triggered averaging, we generated electrical footprints (EFs) for individual neurons. Skeletonization of these EFs allowed us to reconstruct and investigate the propagation paths that action potentials travelled along the neurites (see Methods). Confirming our immunohistochemical data, we found that the inferred EFs indicated unipolar, bipolar, pseudounipolar and multipolar neuron morphologies (Fig. 6F). Such detailed spatial analysis was made possible by the very dense packaging of electrodes on these HD-MEAs, i.e., an electrode-to-electrode pitch of 17.5 µm, and simultaneous high temporal resolution (10 or 20 kHz sampling rate). For further analysis, we only analyzed neurons that had their EF initiation site clearly located on the array. The mean conduction velocity pooled across weeks 4 to 9 was 730.3 ± 25.35 µm/ms (mean ± SEM) (Fig. 6G). We then assessed the mean conduction velocity in the two axon branches separately. Interestingly, our results indicated that EFs in the pseudounipolar class had branches that clearly differed in their action potential propagation speeds, with a mean conduction velocity of 821.1 ± 27.92 µm/ms in the faster and of 636.0 ± 24.3 µm/ms in the slower axon branch, representing a putative peripheral and central axon branch, respectively, with a mean maximum velocity difference between peripheral and central axon branches of 30.7 ± 2.2 % (Fig. 6H-J). This might reflect differences in, e.g., the thickness of their neurites or putative different functional roles (Czéh et al., 1977; Waddell et al., 1989). Successful acquisition of a pseudounipolar morphology, accompanied by the functional asymmetry of the two axon branches by iNBI-SNs co-cultured with rat DRG cells further supports the observation that these neurons exhibit authentic hallmarks of a sensory neuron identity.

## Discussion

Transcription factor-based forward programming presents a powerful alternative to conventional small molecule-guided protocols for the generation of neurons from iPSCs. Our findings demonstrate that inducible overexpression of NGN1, BRN3A and ISLET1 is sufficient to directly induce a sensory neuron fate from human iPSCs. These forward programmed iNBI-SNs recapitulate important hallmarks of primary sensory neurons, including expression of typical marker genes including Nav1.7, Nav1.8 and Nav1.9, the potential to functionally connect to CNS type neurons, and morphological maturation towards a pseudounipolar morphology. Furthermore, they provide a suitable tool for disease-associated pathophenotyping and drug testing.

### Forward programming as tool to recreate master transcription factor-based lineage specification

Forward programming promotes lineage entry through forced initiation of defined transcriptional programs. These approaches enable rapid and efficient neuronal conversion with improved reproducibility and reduced heterogeneity across cell lines (Flitsch et al., 2020; McDaid et al., 2025; Y. Zhang et al., 2013; Zhao et al., 2023). In contrast, conventional differentiation paradigms often rely on the stepwise modulation of developmental signaling pathways, thereby more closely recapitulating embryonic development, at the cost of prolonged culture times and increased variability, including sensitivity to plating density during differentiation (Röderer et al., 2023; Schwartzentruber et al., 2018). Thus, both strategies can be viewed as complementary with TF-based differentiation paradigms offering advantages for a rapid lineage entry, higher control of subtype-specific cellular identity and easy scalability.

While our differentiation protocol is based on forced overexpression of NGN1, BRN3A and ISLET1, we found that NGN1 and BRN3A suffice to induce PRPH+ neuronal cultures, as previously described for fibroblast transdifferentiation (Blanchard et al., 2015). However, inclusion of ISLET1, which is known to act in concert with BRN3A during dorsal root ganglion development (Dykes et al., 2011; Lanier et al., 2009; Sun et al., 2008), resulted in an earlier stop of proliferation and a faster fate entry. This observation is corroborated by an analogous study that reported higher direct transdifferentiation efficiency of fibroblasts into sensory neurons when overexpression of Ascl1 and Brn3a was complemented by Islet1 (Xiao et al., 2020).

### Limitations in fine tuning sensory neuron subtype identity

The micro-network of NGN1, BRN3A and ISLET1 plays a key role during early developmental transition of neural crest cells into sensory neuron subtypes (Meltzer et al., 2021; Vermeiren et al., 2020). These transcription factors act in two developmental waves, where first-wave born neurons are more prone to adopt a large- or intermediate-diameter subtype fate and second-wave neurons a small-diameter identity (Meltzer et al., 2021). Hence, both developmental waves of peripheral sensory neurons are guided by co-expression of the three here used transcription factors (Cranfill & Luo, 2021; Meltzer et al., 2021). This is in line with our observation that iNBI-SNs represent a mixed population rather than a single sensory neuron subtype. Our RNAscope analysis further indicates a progressive diversification of iNBI-SNs over time in culture. Comparison of their single nucleus transcriptomic signature at DIV 63 with an *in vivo* DRG atlas suggests highest similarity with mechanoreceptor and nociceptor sensory neuron subtypes. Yet, the fact that iNBI-SNs represent properties of different sensory neuron subtypes is a limitation for e.g. when studying subtype-specific sensory neuron pathologies. Additional modulation of transcription factors that play a developmental role in subtype-specification, such as PRDM12, RUNX1, RUNX3, SHOX2 and KLF7 might enhance enrichment for specific subtypes of sensory neurons. For example, Hulme et al. employed either RUNX3 or SHOX2 in combination with the neuronal pioneer transcription factor NGN2 in order to generate cultures enriched for proprioceptors or mechanoreceptors, respectively (Hulme et al., 2024). Initiating transcription factor overexpression in established neural crest stem cells instead of undifferentiated iPSCs might represent another route to higher fidelity. Along a similar line, Nickolls et al. have shown a higher degree of specialization into mechanoreceptive neurons when overexpressing NGN2 with BRN3A in iPSC-derived neural crest cells instead of iPSCs (Nickolls et al., 2020). As for the well-established NGN2 system for the generation of central excitatory neurons, it was recently described that expression level and duration of the NGN2 overexpression can affect the differentiation outcome, and that concomitant addition of extrinsic factors and morphogens can be employed to further refine and guide neuronal subtype differentiation (Lin et al., 2025; Lin et al., 2021). An additional approach to refine the forward programming paradigm could be to sequentially activate the employed transcription factors, rather than a simultaneous induction, mimicking the temporally staggered expression of these factors during *in vivo* development.

Yet, it is fair to state that *in vitro* lineage instruction can only be a reductionist mimicry of *in vivo* development, where waves of transcription factors are initiated by interdependent extrinsic factor gradients in distinct temporal sequences, and where differentiating sensory neurons undergo further refinement elicited by their immediate environment and their target tissue. This includes peripheral glial cells within the DRG, neurons of the dorsal horn of the spinal cord as well as e.g. different types of skin and muscle cells in the periphery that eventually instruct further specialization such as skin type-dependent formation of touch end-organs (Koutsioumpa et al., 2023).

### iNBI-SNs exhibit near authentic but incomplete functional maturation

The generated iNBI-SNs are electrophysiologically functional, with 21-day-old neurons already showing the capacity to fire action potentials, and a low percentage of spontaneously active neurons of about 15 % across differentiation. Action potential parameters were found to be similar to other iPSC derived sensory neurons (Eberhardt et al., 2025; Röderer et al., 2023) and largely in the same order of magnitude as for primary human DRG neurons (Lesnak et al., 2026). Moreover, when cultured adjacent to CNS-type NGN neurons, iNBI-SNs established connections and, upon optogenetic stimulation, provided functional input to their CNS counterparts. Importantly, virtually all neurons exhibited tetrodotoxin-resistant sodium currents, a hallmark of peripheral sensory neurons. Typically, the voltage-gated sodium channels Nav1.8 and Nav1.9 are associated with TTX-resistant currents in nociceptive sensory neurons and pain associated channelopathies (Bennett & Woods, 2014). We were able to identify a TTXr current, resembling Nav1.8 and its gating characteristics that was blocked by VX-150, but not a clear slow-inactivating and persistent current generated by Nav1.9, even though expression of the gene was clearly detected in a subset of iNBI-SNs. Enrichment or fluorescent labeling of this subset of neurons might enable measurement of human Nav1.9 currents in future studies. Therefore, the remaining TTX-resistant current detected in iNBI-SNs might be attributed to the fast activating and inactivating Nav1.5 channel, which is encoded by *SCN5A* and expressed in embryonic and to a lesser extent in adult rodent, as well as in human dorsal root ganglion neurons (Bhuiyan et al., 2025; Kerr et al., 2007; Renganathan et al., 2002; C.-J. Zhang et al., 2026). Despite shortcomings with respect to recapitulating the functionality of fully mature adult SNs, iNBI-SNs do recapitulate previously described hyperactivity phenotype of inherited erythromelalgia patient-derived sensory neurons (Alich et al., 2023; Cao et al., 2016; Meents et al., 2019). Furthermore, the functional pathophenotypes were successfully rescued by application of Nav1.7 inhibitors, indicating the suitability of iNBI-SNs for both disease pathophenotype modeling and drug testing.

### Acquisition of a pseudounipolar phenotype as hallmark property of peripheral sensory neurons

PSC-derived SNs in mono-culture, including iNBI-SNs, typically lack the characteristic pseudounipolar morphology observed in peripheral sensory neurons *in vivo*. As previous reports suggested that the interaction of primary sensory neurons with peripheral glial cells is key for inducing a morphological transition towards pseudounipolarity in rodents (Mudge, 1984; Nascimento et al., 2018, 2022), we co-cultured iNBI-SNs with mixed primary embryonic rat DRG cells. Indeed, this co-culture induced in a cell-contact dependent manner pseudounipolarization of iNBI-SNs, underlining the notion that this morphological switch is triggered via a contact-dependent mechanism. In live imaging experiments few pseudounipolarized neurons returned to a bipolar morphology, which might further indicate that the full morphological maturation is a two-step process with initial induction, followed by stabilization. LeBlang et al. traced the process of pseudounipolarization back to a sensory neuron and satellite glial cell interaction, mediated via semaphorin signaling (LeBlang et al., 2024). Under *in vivo* disease conditions pseudounipolar sensory neurons have been observed to sprout and lose their pseudounipolar morphology (Shiers et al., 2024), a transition that might now become accessible to in vitro experimentation. In our cultures we could observe a transition into a morphological asymmetry recapitulating pseudounipolarization and formation of a thin central and a larger peripheral axon. This morphological asymmetry was accompanied by a functional asymmetry indicating a higher conduction velocity in the peripheral axon branch. Furthermore, a controlled co-culture system comprising iNBI-SNs and isolated or PSC-derived satellite glial cells could further facilitate the investigation of cell-cell interactions and signaling pathways underlying morphological maturation and the acquisition of a pseudounipolar morphology.

We expect that this property can be exploited in further studies for differentiating disease-, trauma or toxic damage and pathologies affecting specifically the peripheral or central compartment of sensory neurons and the DRG. Moreover, building on our observation that iNBI-SNs are capable of forming functional connections with CNS-type neurons, such morphologically refined cultures may provide a powerful platform to reconstruct key aspects of peripheral-central connectivity, reenacting the interplay of the peripheral nervous system with the central circuitry of the spinal cord dorsal horn in a controlled and accessible in vitro setting. Such approaches parallel recent 3D assembloid systems that recapitulate signal propagation through several anatomical compartments of the sensory pathway (J. Kim et al., 2025) and offer a promising avenue to systematically dissect circuit-level mechanisms underlying somatosensory processing.

## Methods

### Cell culture and iPSC line engineering

#### Human iPSC lines

Peripheral sensory neurons were generated from the control iPSC lines WT1 (UKBi013-A (https://hpscreg.eu/cell-line/UKBi013-A)), WT2 (UKBi019-A (https://hpscreg.eu/cell-line/UKBi019-A)), as well as inherited erythromelalgia patient derived cell lines IEM1 (RCi001-A (https://ebisc.org/RCi001-A)) and IEM2 (RCi002-A (https://ebisc.org/RCi002-A)). Excitatory iGlut neurons were generated from the control iPSC line UKBi006-A (https://hpscreg.eu/cell-line/UKBi006-A). The use of iPSC lines was approved by the Ethics Committee of the Medical Faculty of the University of Bonn (approval number 275/08), and informed consent was obtained from all donors. iPSCs were cultured as described previously (Röderer et al., 2023).

### Editing of AAVS1-safe harbor locus

The AAVS1-TRE3G-GFP plasmid and the corresponding TALEN pair were kindly provided by Su-Chun Zhang (Addgene plasmids # 52342; # 52341; # 52343) (Qian et al., 2014). To establish the AAVS1-NGN1-BRN3A-ISLET1 targeting construct, the GFP coding sequence was removed from the AAVS1-TRE3G-GFP plasmid by restriction digest with MluI-HF and SalI-HF (R3198S, R3138S, New England Biolabs) according to manufacturer’s instructions and replaced with the respective transcription factor cassette via HiFi assembly (New England Biolabs). Transcription factor cassettes were designed using the NEBuilder Assembly Tool (https://nebuilder.neb.com/#!/). Transcription factors NGN1 and ISLET1 were cloned from Origene DNA plasmids (NM_006161 and NM_002202, respectively) while BRN3A was cloned from the Teto BRN3A plasmid, which was a gift from Kristin Baldwin (Addgene plasmid # 62221)(Blanchard et al., 2015). Plasmids were amplified in NEB5-alpha competent E. coli (New England Biolabs, C2987H9).

AAVS1-targeted iPSC-lines were generated by nucleofecting 2 million cells with 3 µg of the corresponding targeting plasmid (NGN1, NGN1-BRN3A or NGN1-BRN3A-ISLET1) and 0.5 µg of each TALEN plasmid using the Amaxa Cell Line Nucleofector Kit V in combination with program B-023. Nucleofected iPSCs were seeded on Geltrex-coated dishes (180 µg/ml) in StemMACS iPS Brew with 10 µM ROCK inhibitor. Puromycin selection (0.5 µg/ml) was started 48 to 72 hours after nucleofection for ca. 5 days. Automated picking of single clones was performed using the CellCelector System as described previously (Elanzew et al., 2020). Clones were expanded mono-clonally and validated for homozygous transgene integration as described previously (Peitz et al., 2020).

### Production, purification, and characterization of rAAV vectors

We used rAAV-hSyn-tdTomato (Addgene, Watertown, USA, plasmid number #50977) and rAAV-hSyn-ChRmine-mScarlet (Addgene, Watertown, USA, plasmid number #130994) to visualize and subsequently optically stimulate sensory neurons. A helper virus-free, three-plasmid-based strategy was applied to produce rAAV viruses (During et al., 2003). Co-transfection of AAV expression vectors and three AAV helper viruses (pRV, pH21, pFdelta6) was performed in HEK293 cells to generate viral particles of chimeric serotype 1/2. Viral particles were then purified using heparin columns. SDS-PAGE and infection of primary neurons and real-time PCR were used to characterize newly generated viruses.

Briefly, 48 hours after transfection fluorescent HEK293 cells were harvested in 9 ml PBS and spun for 10 min at 800 rpm and 4 °C. Each pellet was resuspended in 45 ml lysis buffer (20 mM Tris-HCl, 150 mM NaOH pH 8,0) and frozen at −20 °C overnight. Upon thawing at RT, 6,6 µl Benzonase (250 – 350 U/µl) and 2,25 ml 10 % NaDOC were added to each tube, to lysate HEK293 cells. Suspensions were incubated at 37 °C for 1 hour and spun for 15 min at 3000 x g and 4 °C. The supernatant was collected and frozen at −20°C for heparin column purification. After thawing at RT, the suspension was centrifuged (15 min, 3000 x g, 4 °C) and the virus-containing supernatant was loaded on a lysis buffer pre-equilibrated 1 ml HiTrap Heparin HP column (SIGMA). Several washing steps later (increasing salt conditions, 100 – 300 mM NaCl), the virus was eluted under high-salt conditions (400 – 500 mM NaCl). Subsequently, the virus was concentrated and rebuffered with PBS in Amicon Ultra tubes and sterile filtered through a 0.2 µm Acrodisc column. Purification and integrity of viral capsid proteins (VP 1-3) were monitored on a coomassie-stained SDS/protein gel. The genomic titers were determined using an AAV quantification Kit (NORGEN Biotek Corporation, Thorold, ON, Canada) in combination with an ABI 7700 real-time PCR cycler (Life Technologies; Carlsbad, California, USA). Viral titers were 3.4 x 10^8^ dsDNA/ml (rAAV hSyn-ChRmine-mScarlet), and 1.74 x 10^8^ dsDNA/ml (rAAV-Syn-tdTomato). rAAV-containing solution was aliquoted in 6 µl aliquots and stored at - 70 °C.

### Differentiation and culture of human iPSC-derived sensory neurons

For forward programming, single cell iPSCs were seeded at a density of 10.000 cells / cm^2^ in StemMACS iPS Brew (Miltenyi Biotec, 130-104-368) with 10 µM ROCK inhibitor (Cell Guidance Systems, SM02) on Geltrex-coated culture dishes (6W-MTP or T175 flask) (180µg/ml). After 24 hours media was switched to Neurobasal Medium (Thermo Fisher Scientific, 21103-049) supplemented with 1% N2 supplement (Thermo Fisher Scientific, 17502-048), 2% B27 supplement (Thermo Fisher Scientific, 17504044), 2 mM (1x) GlutaMAX (Thermo Fisher Scientific, 35050038), 0.02 mM 2-Mercaptoethanol (Thermo Fisher Scientific, 31350010), 12 µg/ml Gentamycin (Thermo Fisher Scientific, 15710049), 200 µM Ascorbic acid (Sigma Aldrich, A4544) and 1µg/ml Doxycyclin (Sigma Aldrich, D9891). Medium was renewed every other day. On day 7 of differentiation sensory neurons were dissociated in Accutase (Thermo Fisher Scientific, A11105-01) and directly seeded for downstream assays or frozen in cold CryoStor CS10 freezing medium (Sigma Aldrich, C2874-100 ML) at −80 °C. Frozen cells were transferred to a liquid nitrogen tank for long-term storage after 24 hours.

The tdTomato-fluorescence or the optogenetic activator channel ChRmine were introduced by AAVs, that were added to the differentiating cells for 48 hours from day 3 to day 5 of differentiation. To prevent cell death during AAV transduction CEPT (Chen et al., 2021) was added to the medium.

Differentiated iPSC-derived sensory neurons were cultured as described before (Röderer et al., 2023). Briefly, iPSC-derived sensory neurons were seeded in Neurobasal A Medium (Thermo Fisher Scientific, 10888022) supplemented with 1% N2 supplement (Thermo Fisher Scientific, 17502-048), 2% B27 supplement (Thermo Fisher Scientific, 17504044), 2 mM (1x) GlutaMAX (Thermo Fisher Scientific, 35050038) and 0.02 mM 2-Mercaptoethanol (Thermo Fisher Scientific, 31350010), 12 µg/ml Gentamycin (Thermo Fisher Scientific, 15710049), 200 µM Ascorbic acid (Sigma Aldrich, A4544), 0.1 mg/ml human recombinant Laminin (BioLamina, LN521), 10 ng/ml GDNF (Cell Guidance Systems, GFH2), 10 ng/ml BDNF (Cell Guidance Systems, GFH1), 10 ng/ml NGF (Peprotech, Cranbury, NJ, 450-01), and 10 ng/ml NT3 (Peprotech, 450-03) supplemented with 10 µM ROCK inhibitor Y-27632 (Cell Guidance Systems, SM02). After 3 days, cells were treated with 1 µg/ml Mitomycin C (Sigma Aldrich, M4287) for 2 hours at 37°C to inactivate remaining proliferative cells. Medium was changed twice per week without ROCK inhibitor.

IPSC-derived sensory neurons differentiated via application of small molecule inhibitors were generated and cultured as previously described (Röderer et al., 2023).

### Rat DRG mixed cultures

DRG cultures were prepared with Wistar rat embryos at the embryonic day 16. All experiments employing primary rodent cells followed the European Union Directive 2010/63/EU and national Decree-law number 113-2013; protocols were approved by the i3S Ethical Committee and the Portuguese Veterinarian Board.

Wistar rat cultures were performed as previously described (Nascimento et al., 2022). Briefly, DRGs were isolated from embryonic day 16 Wistar rat embryos and digested with trypsin 0.05 % EDTA (Gibco, #25300062), for 1 hour at 37°C. Digestion was interrupted with 5% fetal bovine serum (FBS) (Sigma, #F9665) diluted in Neurobasal (Gibco, #21103-049). Later, cells were centrifuged and mechanically dissociated, with a 1000 µl micropipette, in the Neurobasal medium supplemented with 2% B27 (Gibco, #17504044), 2 mM L-glutamine (Gibco, #25030024), 1% penicillin–streptomycin (Gibco, #15140-122), and 50 ng/ml of nerve growth factor (Millipore, #N-100 or Peprotech, 450-01). 25,000 cells were seeded per well in a 24-well plate (Corning, #734-1606), previously coated with 20 μg/ml poly-L-lysine (Sigma-Aldrich, #P2636) and 5 μg/ml laminin (Sigma-Aldrich, #L2020). Medium was partially changed once a week.

### Co-Culture of iNBI-SN with mixed rat DRG cells

INBI-SNs were co-cultured with rat glial cells, via non-filtered media or direct transfer, from DIV 7 onwards. For the conditioned media condition, medium from 8-12 DRG rat wells (approximately 300 µl per well) was collected and completed with the same amount of complete sensory neuron media and filtered with a 0.45 µm filter (Neoreax, #M000070), this combination of rat and sensory neuron media was used to perform media changes on the iNBI-SN mono-cultures. For this, 200 µl of old culture medium was removed and replaced with 300 µl of the mixture. For the co-culture condition, rat glial cells were detached from the coverslip using trypsin 0.05 % EDTA for 15 minutes at 37°C. Detached cells were collected and 10 % FBS was used to stop trypsin activity. Cells were centrifuged (5 min, 200 x g), resuspended in 1 ml of complete sensory neuron media (see Differentiation and culture of human iPSC-derived sensory neurons) and counted. Rat DRG glial cells were added to the iNBI-SN cultures at a ratio of 1:2, i.e. 30,000 glial cells on top of 60,000 iNBI-SN.

### Quantitative real time PCR

Total RNA samples were extracted using the Maxwell RSC simplyRNA tissue kit (Promega, AS1340) following the manufacturer’s protocol. cDNA was generated by reverse transcription using the qScript cDNA synthesis kit (Quantabio, 95047-100), employing 500ng total RNA. Primers used in this study are listed in Table S1, all primer pairs were tested for specificity and linear amplification across a cDNA dilution series prior to experiments. Quantitative real-time PCRs (qRT-PCR) were performed on an Applied Biosystems ViiA7 Real-Time PCR System, using the SYBR Green detection method, with *GAPDH* serving as a houskeeping gene. qRT-PCR results were analyzed with the 2^-ΔΔCt^ method (Livak & Schmittgen, 2001).

### Immunocytochemistry and imaging

Cells were fixed for 10 min with 4 % paraformaldehyde (Electron Microscopy Sciences, #16316 or Thermo Fisher Scientific, J61899). Afterwards, cells were washed with DPBS (Thermo Fisher Scientific, #14040133 or #14190-094) and permeabilized with 0.3% Triton X-100 (Sigma, #T9284) diluted in DPBS. Endogenous autofluorescence was blocked in co-cultures with 0.2M of ammonium chloride (Merck, #1.01145.0500), for 10 minutes at room temperature. Next, cells were blocked with 5% normal donkey serum and (Sigma, #D9663) 0.1% Triton X-100 diluted in DPBS, for 1 hour at room temperature. Samples were incubated with the primary antibodies (see Table S2), diluted in blocking solution, overnight at 4°C. On the following day, cells were incubated with the secondary antibodies (see Table S2), for 1 hour at room temperature. Cell nuclei were stained with DAPI (Bio-Rad, #1351303 or Sigma, #D9542), for 5 minutes. Samples were coverslipped with mounting media (Ibidi, #50001 or Invitrogen, #00-4959-52) and kept at 4°C.

Fluorescent images were acquired with a Leica DMI6000 widefield inverted motorized microscope (Leica Microsystems, Germany), a ZEISS Axio Imager Z1 equipped with an Apotom 1.0 (Zeiss, Germany) or an InCell Analyzer 2200 plate microscope (GE Healthcare, USA).

### RNAscope *in situ* hybridization

RNAscope (multiplex fluorescent v2 kit, Cat 323100) was performed as instructed by Advanced Cell Diagnostics (ACD-bio). The labeling was performed following the “RNAscope Multiplex Fluorescent v2 Assay for Cultured Adherent Cells” protocol. RNAscope probes and fluorescent TSA dyes were used and prepared according to the labeling plan, as shown in Table S3. The coverslips were mounted on glass slides with Aqueous Fluoroshield with DAPI (Abcam, Catalog No.: ab104139). For analysis, eight windows per coverslip were imaged on epifluorescence setup of Zeiss Axio Observer at 20x magnification (Plan-Apochromat 20x/0,8 M27). Representatives were imaged with confocal Zeiss LSM900 at 20x magnification (Plan-Apochromat 20x/0,8 M27) with a Z-stack setup covering 12 µm with a step size of 1 µm, a scanning speed of 7, and 8 px/µm image resolution. Image acquisition and processing were performed on Zen 3.5 (ZEN system) software. Quantification was performed using HALO v3.6.4134.263 and HALO AI 3.6.4134 modules. The “Nuclei Seg (Halo-AI)” machine learning classifier was trained to identify and segment neurons. Quantification of mRNA puncta was done using the module “Indica Labs – FISH-IF v2.2.5”. Visualizations and statistical analysis were performed with GraphPad Prism 9.4.0 (GraphPad Software, Inc. San Diego, CA USA).

### Bulk RNAseq gene expression analysis

RNA was extracted to perform bulk-RNA sequencing, from differentiation cultures at DIV7 and after replating at DIV 35 and 63. RNA was extracted using Agilent Absolutely RNA Microprep reagents (Catalog # 400805). Four technical replicates were generated for every time point. Concentration and total RNA amounts were estimated using Bioanalyzer.

RNA samples from each time point were sent to Novogene (UK) Company Limited, Cambridge, UK for library preparation and sequencing. Messenger RNA was purified from total RNA using poly-T oligo-attached magnetic beads. After fragmentation, the first strand cDNA was synthesized using random hexamer primers, followed by the second strand cDNA synthesis using dTTP for non-directional library. For the non-directional library, it was ready after end repair, A-tailing, adapter ligation, size selection, amplification, and purification. The library was checked with Qubit and real-time PCR for quantification and bioanalyzer for size distribution detection. Quantified libraries were pooled and sequenced on Novaseq X Plus with PE150 strategy RNAseq (Poly A enrichment - 9G data output per sample).

### Data Quality Control

Original image data file from Illumina’s was transformed to raw sequence reads by CASAVA base recognition and stored in FASTQ. Both Sequencing Error Rate and GC Content Distribution were inspected and all sample were considered having high quality (Sequencing Error Rate <1% and constant and stable GC content distribution). Raw reads were then filtered and any reads with adapter contamination, reads with more than 10 percent of uncertain nucleotides and reads with low quality nucleotides (Base Quality less than 5) constituting more than 50 percent of the read, were removed.

### Alignment and Gene Expression Quantification

Alignments were performed with HISAT2 (v.2.0.5, (D. Kim et al., 2019)) to the human genome reference (version hg38). Quantification of gene expression levels was done with featureCounts (v.1.5.0-p3, (Liao et al., 2014)) into FPKM read counts (number of Fragments Per Kilobase of transcript sequence per Millions base pairs sequenced). Gene expression distribution showed no outliers and correlation analysis of samples showed a Pearson’s correlation R^2^>0.9 between all biological replicates.

### Principal Component Analysis

Principal Component Analysis (PCA) was performed on the gene expression value (FPKM) of all samples, using the DESeq2 package (1.20.0, (Love et al., 2014)) in R (R Core Team, 2021).

### Differential gene expression analysis

Differential expression analysis (DEA) was performed using the DESeq2 R package (1.20.0, (Love et al., 2014)). The resulting P-values were adjusted using the Benjamini and Hochberg approach for controlling the false discovery rate (FDR). Genes with an adjusted P-value ≤0.05 found by DESeq2 and absolute foldchange of ≥1 were assigned as differentially expressed. All the differentially expressed genes in the comparison group were pooled as the differential gene set. We used the mainstream hierarchical clustering to cluster the FPKM values of genes and homogenized the row (Z-score). Results were plotted as heat maps, with samples with similar expression patterns gathered together.

### Single-nucleus RNA sequencing

#### Sample preparation

Single-nuclei were isolated from freshly harvested iNBI-SNs in Nuclei EZ lysis buffer (NUC101-1KT, Sigma-Aldrich) by dounce homogenization. After filtering and washing through a 40 µm strainer, nuclei were pelleted (500 x g, 5 min, 4°C), resuspended in wash buffer (1 % BSA w/v, 0.2 U/µl RNase inhibitor in DPBS (D8537, Sigma-Aldrich), and DAPI-positive nuclei were purified by sorting (FACS Aria II, BD). Single nucleus RNA-libraries were generated using the Chromium Next GEM Single Cell 3’ Kit (v3.1, 10x Genomics). Approximately 16,500 cells were added to each reverse transcription mix and loaded on one lane of a microfluidic G chip. Downstream reverse transcription, cDNA synthesis/amplification, and library preparation were performed according to the manufacturer’s instructions. All samples were sequenced at the Cologne Center for Genomics (CCG) (Cologne, Germany) on a NovaSeq 6000 (Illumina) with 2×100 bp sequenced into the 3′ end of the mRNAs. The CCG provided QC metrics and Cell Ranger output files. All gene expression tables were then imported into R Statistical Software (R Core Team, 2021) in RStudio (Posit Team, 2025), merged into one object and analysed with Seurat (v.5.1.0,(Stuart et al., 2019) using standard procedures.

#### Initial quality control, clustering and visualization of snRNAseq

To be included for analysis, nuclei were required to contain counts of greater than 1100 unique genes and 1200 unique molecular identifiers (UMIs), fewer than 5 % of the counts deriving from mitochondrial genes, and a ratio of genes per UMI above 0.8. These criteria were met by 7,240 nuclei.

Next, the data was normalized by Seurat’s global-scaling normalization function *NormalizeData()* with default settings. The top 2,000 variable features were identified and linear transformed with Seurat’s *ScaleData()* function. The most significant principal components (PCs) were identified by elbow plot and the top 30 principal components (PC) were retrieved with *RunPCA()*. To integrate nuclei from both cell lines, anchor-based *CCA integration* was performed as implemented in Seurat. For visualization, a dimensional reduction was performed by calculating the Uniform Manifold Approximation and Projection (UMAP) coordinates in the integrated reduction space, utilizing *runUMAP()*. Based on 30 PCs, clustering was performed with Seurat’s *FindNeighbors()* and *FindClusters()* functions with resolution set to 0.3. Initially, 10 clusters were identified (Suppl. Fig. 4B)

The procedure was repeated for a subset of nuclei, identified as neurons by gene ontology (GO)-Analysis (Suppl. Fig. 4C). The subset was again normalized, scaled and the top 2,000 variable features and the top 30 PCs retrieved as described above. Again, *CCA integration* was performed and a new UMAP calculated. Clustering was performed based on 15 PCs with resolution set to 0.3.

#### Cluster Marker and GO-Analysis

For each cluster, marker genes were identified using the *FindAllMarkers()* function after joining all layers. Markers were then filtered for genes with an average logarithmic fold change greater than 1 (avg_log2FC >1). Those marker lists were then used for GO-Analysis using the *compareCluster()* function of the ClusterProfiler package (Yu et al., 2012). Gene symbols were converted to Entrez IDs using the org.Hs.eg.db package (*Bioconductor - Org.Hs.Eg.Db*, n.d.). P values were adjusted by the Benjamini-Hochberg method and *p* and *q* value cut-offs were set to 0.05.

#### Label Transfer

To approximate the functional identity of the nuclei classified as matured neuron, a label transfer was used, utilizing the human DRG Visium transcriptomics dataset published by Tavares-Ferreira et al. 2022 (Tavares-Ferreira et al., 2022).

The dataset was retrieved from the institutional repository of the University of Texas at Dallas (https://utdallas.app.box.com/s/dgs8a7vo9uu9cx6l81somaxb468hpgnz, last accessed in August 20, 2026), normalized and the most variable features were retrieved as described above. In the next step, transfer anchors were identified using the *FindTransferAnchors*() function with 15 PCs. After, cluster labels were transferred to the query using the *TransferData()* function.

To plot the iNBI-SN dataset and the Tavares dataset in the same UMAP, both datasets were merged and normalized. After retrieving the 20,000 most variable features, the dataset was scaled and the top 30 PCs calculated. Then labels were transferred from the Tavares dataset onto the merged dataset. Using Seurat’s *MapQuery()* function, the UMAP of the single Tavares dataset was transferred to the merged dataset.

#### Visualization

Graphs were generated using functions implemented in the specified packages or with the ggplot2 package (Wickham, 2016).

### Proteomics

#### Sample preparation

Human iPSC-derived sensory neurons were harvested at the indicated time points. Culture media was removed and cells were washed with DPBS (Thermo Fisher Scientific, #14190-094). Cells were scraped from plates and collected in 15ml tubes, plates were washed with 1ml ice cold DPBS and added to the tubes. Samples were kept on ice till centrifugation. After collection of all samples cells were pelleted in a cooled centrifuge at 400 g at 4°C for 5 minutes. Cells were resuspended in a suitable amount of T-PER buffer (ThermoFisher, #78510), supplemented with protease inhibitor (Sigma, P2714), kept on ice for 20 minutes and vortexed every couple of minutes. Cell debris was pelleted at 4°C for 10 minutes at full speed. Supernatant was carefully transferred to Protein LoBind tubes (Eppendorf, 0030108116), frozen and stored at −20°C. Proteome sample concentrations were determined with BCA-Assay (Thermo Scientific, 23227) according to manufacturers’ protocol. Samples were then diluted to 2 μg/μl in T-PER buffer with protease inhibitor.

SDC buffer containing 1% sodium deoxycholate (Sigma-Aldrich), 40 mM 2-chloroacetamide (Sigma-Aldrich), 10 mM tris(2-carboxyethyl)phosphine (Thermo Fisher Scientific) and 100 mM Tris at pH 8.0 was added 1:1 to the lysed samples followed by incubation at room temperature for 20 min. Then, samples were diluted 1:1 with water and digested for 1.5 hours at 37 °C with 1 µg of LysC and overnight at 37°C with 1 µg of LysC and 3 µg of trypsin (Promega). The peptide mixture was acidified with trifluoroacetic acid (Merck) to a final concentration of 1%, followed by desalting of the peptides via SCX StageTips. Samples were vacuum dried and re-suspended in 6 µl of buffer A (0.1% formic acid).

#### LC-MS/MS data acquisition

The peptides were loaded onto a 30-cm column (inner diameter: 75 microns; packed in-house with ReproSil-Pur C18-AQ 1.9-micron beads, Dr. Maisch GmbH) via the autosampler of the Thermo Easy-nLC 1200 (Thermo Fisher Scientific) at 60 °C. Eluting peptides were directly sprayed onto the mass spectrometer Exploris 480 (Thermo Fisher Scientific). Peptides were separated with a flow rate of 250 nL/min by a gradient of buffer B (80% ACN, 0.1% formic acid) from 2% to 30%B over 105 minutes followed an increase to 60%B over 5 minutes then to 95%B over the next 5 minutes and finally the percentage of buffer B was maintained at 95%B for another 5 minutes.

The mass spectrometer was operated in a data-dependent mode with MS1 scans from 300 to 1750 m/z (resolution of 60000 at m/z =200), and up to 15 of the top precursors were selected for fragmentation using higher energy collisional dissociation (HCD with a normalized collision energy of value of 28). The MS2 spectra were recorded at a resolution of 15000 (at m/z = 200).

The AGC (Automatic Gain Control) target for both MS and MS/MS scans was set to 3E6 and 1E5, respectively, with a maximum injection time set to “auto”. Dynamic exclusion was set to 30 seconds.

#### Data Analysis

Raw data were processed using the MaxQuant computational platform (version 2.0.1.0) [https://www.nature.com/articles/nbt.1511]. Shortly, the peak list was searched against the Uniprot database of human (SwissProt) with an allowed precursor mass deviation of 4.5 ppm and an allowed fragment mass deviation of 20 ppm. MaxQuant by default enables individual peptide mass tolerances, which was used in the search. Cysteine carbamidomethylation was set as static modification, and methionine oxidation and N-terminal acetylation as variable modifications. The match-between-run option was enabled, and proteins were quantified across samples using the label-free quantification algorithm in MaxQuant generating label-free quantification (LFQ) intensities.

#### Statistical Analysis

The Perseus software suite (v. 1.6.14.0) (Tyanova et al., 2016) was used to remove protein groups identified only by site, as well as contaminants and reverse hits. Protein groups that were not detected in at least 2 out of 3 replicates for the tested conditions were excluded. The data was then log(2) transformed. Student’s t-tests and volcano plot analysis were conducted, with false discovery rate (FDR) correction applied to control for multiple testing. PCA analysis was also conducted on the protein groups that were present in at least 2 out of 3 replicates for each condition, and the missing values were imputed from a downshifted normal distribution (width 0.3, downshift 1.8 SD, per sample).

#### Gene Ontology Enrichment Analysis

PANTHER was used to run statistical overrepresentation analysis for the GO term biological process (Mi et al., 2019; Thomas et al., 2022). The PANTHER Overrepresentation test was performed in the version released on 2024-08-07 with the GO Ontology database DOI: 10.5281/zenodo.12173881, released on 2024-06-17.

### Electrophysiology

#### Current-clamp recordings

Current-clamp whole-cell recordings of iPSC-derived sensory neurons were performed as previously described in Röderer et al., 2023.

#### Voltage-clamp recordings

Voltage-clamp recordings were performed in two independent laboratories with a slightly different methodology: Voltage clamp whole-cell recordings of iPSC-derived sensory neurons were performed between DIV35 and 70, as indicated in the figure legends. Recordings were conducted at room temperature (∼20°C) using either an EPC 10 USB patch-clamp amplifier (HEKA Electronics, Germany) or a Multiclamp 700B amplifier with pClamp 11 software (Axon Instruments). Data acquisition was performed at a sampling rate of 50 kHz with a low-pass filter set to 10 kHz. The extracellular solution contained either: 140 mM NaCl, 1 mM CaCl_2_, 1 mM MgCl_2_, 10 mM HEPES, 20 mM TEA-Cl, 1 mM 4-Aminopyridine, 0.1 mM CdCl_2_, and 1 mM D-glucose (pH 7.4, 327 mOsm) or 120mM NaCl, 3 mM KCl, 2 mM CaCl_2_, 1 mM MgCl_2_, 10 mM HEPES, 0.1 mM NiCl_2_, 20 mM TEA-Cl, 0.1 mM CdCl_2_ and 10 mM glucose (pH 7.4, 300 mOsm). Intracellular solutions were filled in fire-polished borosilicate glass pipettes containing: 10 mM NaCl, 140 mM CsF, 10 mM HEPES, 1 mM EGTA, 5 mM D-glucose, and 5 mM TEA-Cl (pH 7.3, 316 mOsm) **or** 70 mM CsCl, 70 mM CsF, 0.5 mM CaCl_2_, 3 mM MgCl_2_, 10 mM HEPES, 5 mM EGTA, 2 mM MgATP, 0.3 mM GTP tris (pH 7.3, 290 mOsm). An IV curve was run after establishing the whole-cell configuration before and after applying pharmacological treatments. A gravity perfusion system was used to apply different pharmacological compounds targeting the Nav channel isoforms expressed in iNBI-SNs. The neurons were first recorded in the ECS with 0.2% DMSO until a stable Nav current amplitude, defined as the Nav current baseline, was established for at least 5 consecutive sweeps at Vmax. Following this baseline, 100 nM SSCI-2 pharmacological blocker (Grünenthal) was applied to determine the percentage of sodium current generated by the sodium channel Nav1.7. Once a stable current amplitude was detected, a combination of 100 nM SSCI-2 + 3 mM VX-150 (Grünenthal) mix was applied to determine the Nav1.8 current component. Finally, an application of 100 nM SSCI-2 + 3 mM VX-150 + 500 nM tetrodotoxin citrate (TTX citrate, Hello Bio) determined the ratio of TTX-sensitive versus TTX-resistant Nav currents in the hiPSC-SNs.

#### Multi-electrode array recordings

24-well multi-electrode array plates (Axion BioSystems, M384-tMEA-24 W) were coated with 0.07% polyethyleneimine (Sigma Aldrich, P3143-100ML) diluted in 1x borate buffer (Thermo Fisher, 28341) and 10 µg/ml human recombinant laminin (Biolamina, LN521) diluted in DPBS containing calcium and magnesium (Thermo Fisher, 14040133). INBI-SNs were seeded at a density of 60,000 cells per well. A Maestro EDGE system, equipped with a Lumos 24 optical stimulation system was used for signal recording and stimulation, controlled by the Axis Navigator software (Axion BioSystems). Raw data were recorded under standard settings with the sampling rate at 12.5 kHz, bandpass between 0.1 and 2000 Hz. Online spike detection was performed with a threshold of 6 times standard deviations. For recordings of spontaneous activity, plates were loaded and equilibrated for 10-15 minutes, followed by 15 minutes recording. For pharmacological experiments, compounds were added at a ratio of 1:10 to reach working concentration. Selective Nav1.7 blocker SSCI-2 was used to assess the effect on sodium channels at DIV64 of maturation at 100 and 300 nM. After the addition of the blocker the plate was recorded for 20 min. Initial baseline recordings and post drug application (5 min) were used to calculate % change in the Mean firing rate and hence the inhibition. Drugs were added against the wall of the well with a dosing volume of 50 μl/well. % change in the mean firing rate was calculated using the Neural metric tool (Axion Biosystems) and graphs were plotted using the Axis Metric Plotting tool and Graph pad prism software. Raster plots were outputted from AxIS Spike files using the Neural Metric Tool software. 10 mM stocks of SSCI-2 were prepared in DMSO and further diluted stepwise to prepare 10X compound concentration in medium. Total well volume was kept at 500 µl and 0.1% DMSO was used as vehicle control. TTX (Tetrodotoxin), 500 nM was applied at the end of experiment. Application of 200 nM or 1000 nM of the Nav1.7 blocker PF-05089771 (Tocris, 5931) was employed to investigate the effect of Nav1.7 inhibition on inherited erythromelalgia patient derived iNBI-SNs.

To assess formation of functional connectivity of iNBI-SNs with CNS neurons, iNBI-SNs were plated spatially separated by a silicon spacer (IBIDI, 80209) together with induced excitatory glutamatergic neurons (Peitz et al., 2020) on to a 6-well multi-electrode array plate (Axion BioSystems, M384-tMEA-6W). Three days after plating the silicon spacer was removed to allow neurite outgrowth and formation of synaptic connections between both populations. Optical stimulation was started after a 5-minute baseline recording, cultures were stimulated 10 times every 30 seconds with a single flash of green (530 nm) light, with 60% intensity, lasting 50 ms. Data was retrieved from iNBI-SN and iGlut covered electrodes separately to investigate signal transmission.

#### High-density micro-electrode array recordings

Custom single-well high-density microelectrode arrays (HD-MEAs), and commercially available multi-well HD-MEA plates (MaxWell Biosystems, Zurich, Switzerland; MaxOne Chip, MaxTwo 6-Well Plates, MaxTwo 24-Well Plates) were filled with complete sensory neuron culture medium to precondition the plates for 2 hours at 37°C. Afterwards, sensory neuron medium was removed and plates were coated with 0.07% PEI (Sigma-Aldrich, P3143) in 1 x borate buffer (Thermo Fisher Scientific, 28341) for 1 hour at 37°C, followed by washing three times with DPBS containing calcium and magnesium (Thermo Fisher Scientific, 14040133). Plates were then left to dry for 30 minutes, coated with 10 µg/ml human recombinant laminin (BioLamina, LN521), and incubated for 2 hours at 37°C. 50,000 iNBI-SN (tdTomato-labeled or unlabeled) were seeded in a volume of 50 µl on to the electrode area, and after 45 minutes plates were filled with complete sensory neuron medium supplemented with 10 µM ROCK inhibitor Y-27632 (Cell Guidance Systems, SM02). INBI-SNs were treated with 1 µg/ml Mitomycin C for 2 hours at 37°C three days after seeding. 25,000 rat DRG cells were added to the cultures at DIV 7, as described previously. Weekly electrophysiological recordings started one week after the plating. Recordings were performed on HD-MEAs comprising a sensing area of 3.85 × 2.10 mm^2^, 26’400 electrodes (of which up to 1024 electrodes can be simultaneously recorded) and 17.5 μm electrode pitch; and a sampling rate of 20kHz (see (Müller et al., 2015) for more details); the MaxTwo multi-well system by MaxWell Biosystems has very similar chip design specifications, but records at a lower sampling rate (10 kHz).

The recording protocol started with the MaxLab Live (MaxWell Biosystems) ActivityScan assay (whole-array scan, 60 seconds/config), followed by an AxonTracking assay (up to 30 hotspots per array, which were selected based on the activity results of the full ActivityScan assay; 120 seconds/config). Default values of the MaxLab AxonTracking assay spike-sorting routine were used to obtain spike-triggered electrical footprints for the subsequent inference of the neuronal conduction velocity of individual neurons.

#### Neuronal conduction velocity estimation

To estimate neuronal conduction velocities from HD-MEA recordings, respectively the electrical footprints of neurons obtained by spike-triggered averaging, involves calculating the velocity of each inferred axonal segment along the paths in the extracted neuronal skeleton. From these segment velocities, the total average conduction velocity for each path was determined, as well as the latency of signal propagation along the entire path.

*Path segment velocity computation* For each path, the velocity of a segment, defined by two directly connected centroids, was computed using the Euclidean distance between the centroids in pixels, converted to micrometers (µm) using a pixel-to-micrometer conversion factor (*px_to_µm*). This conversion factor was derived by mapping the pixel dimensions of the GIF (908 x 497 pixels) to the physical dimensions of the HD-MEA electrode array (3.85 x 2.10 mm²). Let *t* be the time per frame in milliseconds (ms). The value of *t* depends on the sampling rate used during the recording, i.e., for 6- or 24-well plates sampled at 10 kHz: *t* = 0.05 ms. For single-well HD-MEAs sampled at 20 kHz: *t* = 0.025 ms. The time interval for each segment was determined by the difference in the frames in which the centroids were detected. The segment velocity (*Vsegment*) was then calculated as:

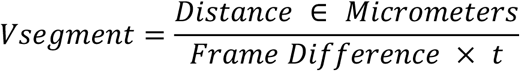

This segment velocity captures the conduction speed of the neuronal signal between two directly connected centroids in a path.

*Total path average velocity computation* The average velocity for an entire path (*Vaverage*) is determined by dividing the total path length (sum of segment distances) by the total time taken to traverse the path. The total time is calculated from the difference between the frame of the last centroid and the frame of the first centroid in the path:

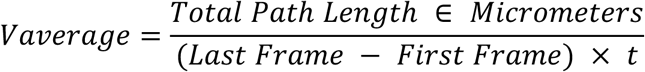

This average velocity captures the average conduction speed of the neuronal signal along the entire path.

*Path latency computation* The latency represents the time delay as the signal travels along the path away from its initiation site. For each centroid i along the path, the latency (*Latencyi*) is calculated as:[]

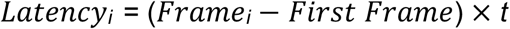

where *t* is again chosen depending on the plate type and sampling rate, as described above. This latency is computed for each centroid, which allows inferring the temporal progression of neuronal activity along the entire path. Further implementation details on the skeletonization and conduction velocity computation process are provided at: https://github.com/stuegerm/BSCTHESIS.

#### Live cell imaging

Morphological changes of differentiating iNBI-SNs and of tdTomato-labeled iNBI-SNs in co-culture with rat DRG cells were monitored using a live cell microscope (Leica Microsystems, DMI6000 B), equipped with a camera (Hamamatsu, Orca-Flash 4.0), CO_2_ controller (Pecon, CO_2_-Controller 2000) and a temperature control unit (Pecon, TempController 2000-2). The cells were kept in a humid chamber at 37°C and 5% CO_2_ during the imaging. Images were taken every two hours.

### Statistics

Statistical analyses were performed using GraphPad Prism 10 (GraphPad Software). Normal distribution of data was tested using the Shapiro-Wilk test. Subsequently performed statistical tests with respective post hoc tests and appropriate corrections for multiple comparisons were performed as indicated in the figure legends. P ≤ 0.05 was considered statistically significant. Significance was reported as * for p ≤ 0.05, ** for p ≤ 0.01, *** for p ≤ 0.001 or **** for p ≤ 0.0001.

## Resource availability

Proteome, bulk and single nucleus gene expression data will be openly accessible after peer-reviewed publication.

Parts of this study have been reported in the PhD thesis ‘Human iPSC-derived in vitro models of the peripheral nervous system for cellular pain and somatosensation research’ by Pascal Röderer (Röderer, 2022).

## Acknowledgement

We thank Michaela Segschneider, Vanessa Frickel and Cornelia Thiele for excellent technical support. We thank Meray Serdar from the University Hospital Essen, Germany for her support of our study.

PR was supported by the BONFOR program of the Medical Faculty of the University of Bonn and the Ministry of Culture and Science of the State of North Rhine-Westphalia under the Stammzell-InTraNetz.NRW project “Authentic-SN”. ALa has received funding from the DFG, German Research Foundation 363055819/GRK2415, DFG, German Research Foundation 368482240/GRK2416, DFG, German Research Foundation LA 2740/6-1. This work was supported by the Flow Cytometry Facility of the Interdisciplinary Center for Clinical Research (IZKF) within the Faculty of Medicine at RWTH Aachen University (RRID:SCR_028757) and the Cologne Center for Genomics (CCG) as part of the West German Sequencing Center (WGGC) by a NGS DFG SEQ3012 funding (KU 1587/10-1). We thank the Mass Spectrometry Core Facility lab at the Max Planck Institute of Biochemistry for their technical support and processing of our proteomics samples.

## Author contributions

Conceptualization, PR and OB; Methodology, PR, ANI and OB; Formal analysis, PR, ACC, ML, FR, FB, LH, JA, OM, HTJ, KLC, AKK, AZ, YA, MS, NH and MSC; Investigation, PR, ACC, ML, FR, FB, LH, JA, AN, OM, HTJ, KLC, AKK, RB, AZ, TB, MS, NH and MSC; Resources, MKS, NCG, NH, MSC, AL and MSO; Writing – original draft, PR and OB; Writing – review & editing, PR, ACC, ML, FR, FB, LH, JA, AN, OM, HTJ, KLC, AKK, RB, VA, AZ, YM, TB, MS, MKS, ANI, NCG, NH, MSC, AL, MSO and OB; Visualization, PR, ACC, ML, OM, HTJ, KLC, YM and MS; Supervision, PR, ANI, VA, NCG, NH, MSC, AL, MSO and OB; Project administration, PR and OB; Funding acquisition, PR and OB.

## References

Abraira, V. E., & Ginty, D. D. (2013). The Sensory Neurons of Touch. Neuron, 79(4), 618–639. 10.1016/J.NEURON.2013.07.051

Alich, T. C., Röderer, P., Szalontai, B., Golcuk, K., Tariq, S., Peitz, M., Brüstle, O., & Mody, I. (2023). Bringing to light the physiological and pathological firing patterns of human induced pluripotent stem cell-derived neurons using optical recordings. Frontiers in Cellular Neuroscience, 16, 676. 10.3389/FNCEL.2022.1039957

Anderson, D. J. (1999). Lineages and transcription factors in the specification of vertebrate primary sensory neurons. In Current Opinion in Neurobiology (Vol. 9, Number 5, pp. 517–524). Current Biology Ltd. 10.1016/S0959-4388(99)00015-X

Bennett, D. L., Clark, X. A. J., Huang, J., Waxman, S. G., & Dib-Hajj, S. D. (2019). The role of voltage-gated sodium channels in pain signaling. Physiological Reviews, 99(2), 1079–1151. 10.1152/physrev.00052.2017

Bennett, D. L. H., & Woods, C. G. (2014). Painful and painless channelopathies. The Lancet Neurology, 13(6), 587–599. 10.1016/S1474-4422(14)70024-9

Bhuiyan, S. A., Nagi, S. S., Sankaranarayanan, I., Semizoglou, E., Usoskin, D., Yang, L., Yu, H., Arendt-Tranholm, A., Bertels, Z., Bhatia, P., Bouchatta, O., Boyer, K., Cervantes, A., Chalif, J., Chintalapudi, H., Cicalo, A., Copits, B., Cronin, C., Curatolo, M.,…Network, N. P. H. P. (2025). A Reference Atlas of the Human Dorsal Root Ganglion. BioRxiv, 2025.11.05.686654. 10.1101/2025.11.05.686654

Bioconductor - org.Hs.eg.db. (n.d.). 10.18129/B9.bioc.org.Hs.eg.db

Blanchard, J. W., Eade, K. T., Szůcs, A., Lo Sardo, V., Tsunemoto, R. K., Williams, D., Sanna, P. P., & Baldwin, K. K. (2015). Selective conversion of fibroblasts into peripheral sensory neurons. Nature Neuroscience, 18(1), 25–35. 10.1038/nn.3887

Cao, L., McDonne, A., Nitzsche, A., Alexandrou, A., Saintot, P. P., Loucif, A. J. C., Brown, A. R., Young, G., Mis, M., Randal, A., Waxman, S. G., Stanley, P., Kirby, S., Tarabar, S., Gutteridge, A., Butt, R., McKernan, R. M., Whiting, P., Ali, Z.,…Stevensl, E. B. (2016). Pharmacological reversal of a pain phenotype in iPSC-derived sensory neurons and patients with inherited erythromelalgia. Science Translational Medicine, 8(335). 10.1126/scitranslmed.aad7653

Chambers, S. M., Fasano, C. A., Papapetrou, E. P., Tomishima, M., Sadelain, M., & Studer, L. (2009). Highly efficient neural conversion of human ES and iPS cells by dual inhibition of SMAD signaling. Nature Biotechnology 2009 27:3, 27(3), 275–280. 10.1038/nbt.1529

Chambers, S. M., Qi, Y., Mica, Y., Lee, G., Zhang, X. J., Niu, L., Bilsland, J., Cao, L., Stevens, E., Whiting, P., Shi, S. H., & Studer, L. (2012). Combined small-molecule inhibition accelerates developmental timing and converts human pluripotent stem cells into nociceptors. Nature Biotechnology, 30(7), 715–720. 10.1038/nbt.2249

Chang, W., Berta, T., Kim, Y. H., Lee, S., Lee, S. Y., & Ji, R. R. (2018). Expression and role of voltage-gated sodium channels in human dorsal root ganglion neurons with special focus on Nav1.7, species differences, and regulation by paclitaxel. Neuroscience Bulletin, 34(1), 4–12. 10.1007/s12264-017-0132-3

Chen, Y., Tristan, C. A., Chen, L., Jovanovic, V. M., Malley, C., Chu, P. H., Ryu, S., Deng, T., Ormanoglu, P., Tao, D., Fang, Y., Slamecka, J., Hong, H., LeClair, C. A., Michael, S., Austin, C. P., Simeonov, A., & Singeç, I. (2021). A versatile polypharmacology platform promotes cytoprotection and viability of human pluripotent and differentiated cells. Nature Methods 2021 18:5, 18(5), 528–541. 10.1038/s41592-021-01126-2

Chiu, I. M., Barrett, L. B., Williams, E. K., Strochlic, D. E., Lee, S., Weyer, A. D., Lou, S., Bryman, G. S., Roberson, D. P., Ghasemlou, N., Piccoli, C., Ahat, E., Wang, V., Cobos, E. J., Stucky, C. L., Ma, Q., Liberles, S. D., & Woolf, C. J. (2014). Transcriptional profiling at whole population and single cell levels reveals somatosensory neuron molecular diversity. ELife, 3. 10.7554/eLife.04660

Costa, A. C., Murillo, B. R., Bessa, R., Ribeiro, R., Silva, T. F. da, Porfírio-Rodrigues, P., Martins, G. G., Brites, P., Kneussel, M., Misgeld, T., Brill, M. S., & Sousa, M. M. (2024). Axon-specific microtubule regulation drives asymmetric regeneration of sensory neuron axons. BioRxiv, 2024.10.10.617525. 10.1101/2024.10.10.617525

Cranfill, S. L., & Luo, W. (2021). The Development of Somatosensory Neurons: Insights into Pain and Itch. Current Topics in Developmental Biology, 142, 443. 10.1016/BS.CTDB.2020.10.005

Czéh, G., Kudo, N., & Kuno, M. (1977). Membrane properties and conduction velocity in sensory neurones following central or peripheral axotomy. The Journal of Physiology, 270(1), 165–180. 10.1113/JPHYSIOL.1977.SP011944

Devor, M. (1999). Unexplained peculiarities of the dorsal root ganglion. Pain, 82(SUPPL.1). 10.1016/S0304-3959(99)00135-9

Dykes, I. M., Tempest, L., Lee, S. I., & Turner, E. E. (2011). Brn3a and Islet1 act epistatically to regulate the gene expression program of sensory differentiation. Journal of Neuroscience, 31(27), 9789–9799. 10.1523/JNEUROSCI.0901-11.2011

Eberhardt, E., Namer, B., Neureiter, A., Körner, J., Jørum, E., Kurth, I., Winner, B., & Lampert, A. (2025). Spontaneous activity in pain patient stem cell–derived sensory neurons arises from one functional subclass. Pain. 10.1097/J.PAIN.0000000000003865

Elanzew, A., Nießing, B., Langendoerfer, D., Rippel, O., Piotrowski, T., Schenk, F., Kulik, M., Peitz, M., Breitkreuz, Y., Jung, S., Wanek, P., Stappert, L., Schmitt, R. H., Haupt, S., Zenke, M., König, N., & Brüstle, O. (2020). The StemCellFactory: A modular system integration for automated generation and expansion of human induced pluripotent stem cells. Frontiers in Bioengineering and Biotechnology, 0, 1155. 10.3389/FBIOE.2020.580352

Flitsch, L. J., Laupman, K. E., & Brüstle, O. (2020). Transcription Factor-Based Fate Specification and Forward Programming for Neural Regeneration. In Frontiers in Cellular Neuroscience (Vol. 14). Frontiers Media S.A. 10.3389/fncel.2020.00121

Galiakberova, A., Ivanov, S., Golov, A., Artyuhov, A., Zolkin, A., Kondratyev, N., Lagunin, A., Golimbet, V., & Dashinimaev, E. (2025). Transcriptomic profiling of neural cultures from the KYOU iPSC line via alternative differentiation protocols. Frontiers in Molecular Neuroscience, 18, 1661986. 10.3389/FNMOL.2025.1661986/BIBTEX

Hulme, A. J., Finol-Urdaneta, R. K., McArthur, J. R., Marzano, N. R., Maksour, S., Thind, A., Guo, Y., Kaul, D., Maddock, M., Friedrich, O., Martinac, B., Adams, D. J., & Dottori, M. (2024). Distinct molecular and functional properties of human induced-proprioceptor and low-threshold mechanoreceptor neurons. BioRxiv, 2024.05.12.593730. 10.1101/2024.05.12.593730

Jung, M., Dourado, M., Maksymetz, J., Jacobson, A., Laufer, B. I., Baca, M., Foreman, O., Hackos, D. H., Riol-Blanco, L., & Kaminker, J. S. (2023). Cross-species transcriptomic atlas of dorsal root ganglia reveals species-specific programs for sensory function. Nature Communications 2023 14:1, 14(1), 1–15. 10.1038/s41467-023-36014-0

Kalia, A. K., Rösseler, C., Granja-Vazquez, R., Ahmad, A., Pancrazio, J. J., Neureiter, A., Zhang, M., Sauter, D., Vetter, I., Andersson, A., Dussor, G., Price, T. J., Kolber, B. J., Truong, V., Walsh, P., & Lampert, A. (2024). How to differentiate induced pluripotent stem cells into sensory neurons for disease modelling: a functional assessment. Stem Cell Research & Therapy 2024 15:1, 15(1), 99-. 10.1186/S13287-024-03696-2

Kerr, N. C. H., Gao, Z., Holmes, F. E., Hobson, S. A., Hancox, J. C., Wynick, D., & James, A. F. (2007). The sodium channel Nav1.5a is the predominant isoform expressed in adult mouse dorsal root ganglia and exhibits distinct inactivation properties from the full-length Nav1.5 channel. Molecular and Cellular Neurosciences, 35(2), 283. 10.1016/J.MCN.2007.03.002

Kim, D., Paggi, J. M., Park, C., Bennett, C., & Salzberg, S. L. (2019). Graph-based genome alignment and genotyping with HISAT2 and HISAT-genotype. Nature Biotechnology 2019 37:8, 37(8), 907–915. 10.1038/s41587-019-0201-4

Kim, J., Imaizumi, K., Jurjuț, O., Kelley, K. W., Wang, D., Thete, M. V., Hudacova, Z., Amin, N. D., Levy, R. J., Scherrer, G., & Pașca, S. P. (2025). Human assembloid model of the ascending neural sensory pathway. Nature 2025, 1–11. 10.1038/s41586-025-08808-3

Körner, J., Howard, D., Solinski, H. J., Moreno, M. M., Haag, N., Fiebig, A., Maxion, A., Bhuiyan, S. A., Toklucu, I., Bott, R. A., Sankaranarayanan, I., Tavares-Ferreira, D., Shiers, S., Inturi, N. N., Eberhardt, E., Ernst, L., Bonaguro, L., Schulte-Schrepping, J., Beyer, M. D.,…Lampert, A. (2026). Molecular architecture of human dermal sleeping nociceptors. Cell, 0(0). 10.1016/J.CELL.2025.12.048

Koutsioumpa, C., Santiago, C., Jacobs, K., Lehnert, B. P., Barrera, V., Hutchinson, J. N., Schmelyun, D., Lehoczky, J. A., Paul, D. L., & Ginty, D. D. (2023). Skin-type-dependent development of murine mechanosensory neurons. Developmental Cell. 10.1016/J.DEVCEL.2023.07.020

Lallemend, F., & Ernfors, P. (2012). Molecular interactions underlying the specification of sensory neurons. Trends in Neurosciences, 35(6), 373–381. 10.1016/j.tins.2012.03.006

Lampert, A., Bennett, D. L., McDermott, L. A., Neureiter, A., Eberhardt, E., Winner, B., & Zenke, M. (2020). Human sensory neurons derived from pluripotent stem cells for disease modelling and personalized medicine. Neurobiology of Pain, 100055. 10.1016/j.ynpai.2020.100055

Lanier, J., Dykes, I. M., Nissen, S., Eng, S. R., & Turner, E. E. (2009). Brn3a regulates the transition from neurogenesis to terminal differentiation and represses non-neural gene expression in the trigeminal ganglion. Developmental Dynamics, 238(12), 3065–3079. 10.1002/dvdy.22145

LeBlang, C. J., Pazyra-Murphy, M. F., Silagi, E., Dasgupta, S., Tsolias, M., Miller, T., Petrova, V., Zhen, S., Gerrish, K., Jovanovic, V., Castellano, D., Ormanoglu, P., Tristan, C., Woolf, C. J., Tasdemir-Yilmaz, O., & Segal, R. A. (2024). Satellite glial contact enhances differentiation and maturation of human induced sensory neurons. BioRxiv, 2024.07.24.604966. 10.1101/2024.07.24.604966

Lee, G., Chambers, S. M., Tomishima, M. J., & Studer, L. (2010). Derivation of neural crest cells from human pluripotent stem cells. Nature Protocols, 5(4), 688–701. 10.1038/nprot.2010.35

Lesnak, J. B., Schaub, M. K., Gomez, K., Calderon-Rivera, A., Loya-Lopez, S., Stewart, R., Jo, S., Fujita, A., Osorno, T., Mydugolam, H., Desai, M., Natarajan, K., Schackmuth, M. K., Moreno, M. M., Shiers, S. I., Cervantes, A., Funk, G., Horton, P., Vines, E.,…Price, T. J. (2026). An improved preservation method for human dorsal root ganglion neurons enables wider access to human molecular pain neuroscience. Cell Reports Methods, 6(5), 101412. 10.1016/J.CRMETH.2026.101412

Li, C. L., Li, K. C., Wu, D., Chen, Y., Luo, H., Zhao, J. R., Wang, S. S., Sun, M. M., Lu, Y. J., Zhong, Y. Q., Hu, X. Y., Hou, R., Zhou, B. B., Bao, L., Xiao, H. S., & Zhang, X. (2016). Somatosensory neuron types identified by high-coverage single-cell RNA-sequencing and functional heterogeneity. Cell Research, 26(1), 83–102. 10.1038/cr.2015.149

Liao, Y., Smyth, G. K., & Shi, W. (2014). featureCounts: an efficient general purpose program for assigning sequence reads to genomic features. Bioinformatics, 30(7), 923–930. 10.1093/BIOINFORMATICS/BTT656

Lin, H. C., Janssens, J., Eisinger, B., Hornauer, P., Kroell, A. S., Santel, M., Pascual-Garcia, M., Okamoto, R., Karava, K., He, Z., Priouret, M., Schröter, M., Camp, J. G., & Treutlein, B. (2025). Human neuron subtype programming via single-cell transcriptome-coupled patterning screens. Science, 389(6756). 10.1126/SCIENCE.ADN6121/SUPPL_FILE/SCIENCE.ADN6121_MOVIE S_S1_TO_S3.ZIP

Lin, H.-C., He, Z., Ebert, S., Schörnig, M., Santel, M., Nikolova, M. T., Weigert, A., Hevers, W., Kasri, N. N., Taverna, E., Camp, J. G., & Treutlein, B. (2021). NGN2 induces diverse neuron types from human pluripotency. Stem Cell Reports. 10.1016/J.STEMCR.2021.07.006

Livak, K. J., & Schmittgen, T. D. (2001). Analysis of relative gene expression data using real-time quantitative PCR and the 2-ΔΔCT method. Methods, 25(4), 402–408. 10.1006/meth.2001.1262

Love, M. I., Huber, W., & Anders, S. (2014). Moderated estimation of fold change and dispersion for RNA-seq data with DESeq2. Genome Biology, 15(12), 550-. 10.1186/S13059-014-0550-8/FIGURES/9

Ma, Q., Fode, C., Guillemot, F., & Anderson, D. J. (1999). Neurogenin1 and neurogenin2 control two distinct waves of neurogenesis in developing dorsal root ganglia. Genes & Development, 13(13), 1717–1728. 10.1101/gad.13.13.1717

Marmigère, F., & Carroll, P. (2014). Neurotrophin signalling and transcription programmes interactions in the development of somatosensory neurons. Handbook of Experimental Pharmacology, 220, 329–353. 10.1007/978-3-642-45106-5_13

Matsuda, S., Kobayashi, N., Wakisaka, H., Saito, S., Saito, K., Miyawaki, K., Momhwki, K., Shigemoto, K., Murakami, S., & Fujiwara, T. (2000). Morphological transformation of sensory ganglion neurons and satellite cells. Biomedical Reviews, 11(0), 39–52. 10.14748/BMR.V11.129

McDaid, G., Vanek, J., Cromer, B., & Sumer, H. (2025). Transcription Factor-Based Differentiation of Pluripotent Stem Cells: Overcoming the Traps of Random Neuronal Fate. Biomedicines 2025, Vol. 13, Page 2783, 13(11), 2783. 10.3390/BIOMEDICINES13112783

Meents, J. E., Bressan, E., Sontag, S., Foerster, A., Hautvast, P., Rösseler, C., Hampl, M., Schüler, H., Goetzke, R., Chi Le, T. K., Kleggetveit, I. P., Le Cann, K., Kerth, C., Rush, A. M., Rogers, M., Kohl, Z., Schmelz, M., Wagner, W., Jørum, E.,…Lampert, A. (2019). The role of Nav1.7 in human nociceptors. PAIN. 10.1097/j.pain.0000000000001511

Meltzer, S., Santiago, C., Sharma, N., & Ginty, D. D. (2021). The cellular and molecular basis of somatosensory neuron development. Neuron. 10.1016/J.NEURON.2021.09.004

Mi, H., Muruganujan, A., Huang, X., Ebert, D., Mills, C., Guo, X., & Thomas, P. D. (2019). Protocol Update for large-scale genome and gene function analysis with the PANTHER classification system (v.14.0). Nature Protocols 2019 14:3, 14(3), 703–721. 10.1038/s41596-019-0128-8

Mudge, A. W. (1984). Schwann cells induce morphological transformation of sensory neurones in vitro. Nature 1984 309:5966, 309(5966), 367–369. 10.1038/309367a0

Müller, J., Ballini, M., Livi, P., Chen, Y., Radivojevic, M., Shadmani, A., Viswam, V., Jones, I. L., Fiscella, M., Diggelmann, R., Stettler, A., Frey, U., Bakkum, D. J., & Hierlemann, A. (2015). High-resolution CMOS MEA platform to study neurons at subcellular, cellular, and network levels. Lab on a Chip, 15(13), 2767–2780. 10.1039/C5LC00133A

Nascimento, A. I., Da Silva, T. F., Fernandes, E. C., Luz, L. L., Mar, F. M., Safronov, B. V., & Sousa, M. M. (2022). Sensory neurons have an axon initial segment that initiates spontaneous activity in neuropathic pain. Brain, 145(5), 1632–1640. 10.1093/BRAIN/AWAC078

Nascimento, A. I., Mar, F. M., & Sousa, M. M. (2018). The intriguing nature of dorsal root ganglion neurons: Linking structure with polarity and function. In Progress in Neurobiology (Vol. 168, pp. 86–103). Elsevier Ltd. 10.1016/j.pneurobio.2018.05.002

Nguyen, M. Q., von Buchholtz, L. J., Reker, A. N., Ryba, N. J., & Davidson, S. (2021). Single-nucleus transcriptomic analysis of human dorsal root ganglion neurons. ELife, 10. 10.7554/ELIFE.71752

Nickolls, A. R., Lee, M. M., Espinoza, D. F., Szczot, M., Lam, R. M., Wang, Q., Beers, J., Zou, J., Nguyen, M. Q., Solinski, H. J., AlJanahi, A. A., Johnson, K. R., Ward, M. E., Chesler, A. T., & Bönnemann, C. G. (2020). Transcriptional programming of human mechanosensory neuron subtypes from pluripotent stem cells. Cell Reports, 30(3), 932–946.e7. 10.1016/j.celrep.2019.12.062

Peitz, M., Krutenko, T., & Brüstle, O. (2020). Protocol for the Standardized Generation of Forward Programmed Cryopreservable Excitatory and Inhibitory Forebrain Neurons. STAR Protocols, 1(1), 100038. 10.1016/j.xpro.2020.100038

Posit Team. (2025). RStudio: Integrated Development Environment for R. Posit Software, PBC, Boston, MA. URL Http://Www.Posit.Co/.

Qian, K., Huang, C. T.-L., Huang, C.-L., Chen, H., Blackbourn, L. W., Chen, Y., Cao, J., Yao, L., Sauvey, C., Du, Z., & Zhang, S.-C. (2014). A simple and efficient system for regulating gene expression in human pluripotent stem cells and derivatives. Stem Cells (Dayton, Ohio), 32(5), 1230–1238. 10.1002/stem.1653

R Core Team. (2021). R: A language and environment for statistical computing. R Foundation for Statistical Computing, Vienna, Austria. URL Https://Www.R-Project.Org/.

Renganathan, M., Dib-Hajj, S., & Waxman, S. G. (2002). Nav1.5 underlies the ‘third TTX-R sodium current’ in rat small DRG neurons. Molecular Brain Research, 106(1–2), 70–82. 10.1016/S0169-328X(02)00411-4

Rhee, H. J., Shaib, A. H., Rehbach, K., Lee, C., Seif, P., Thomas, C., Gideons, E., Guenther, A., Krutenko, T., Hebisch, M., Peitz, M., Brose, N., Brüstle, O., & Rhee, J. S. (2019). An Autaptic Culture System for Standardized Analyses of iPSC-Derived Human Neurons. Cell Reports, 27(7), 2212–2228.e7. 10.1016/J.CELREP.2019.04.059

Röderer, P. (2022). Human iPSC-derived in vitro models of the peripheral nervous system for cellular pain and somatosensation research. PhD Thesis, University of Bonn. https://bonndoc.ulb.uni-bonn.de/xmlui/handle/20.500.11811/9532

Röderer, P., Belu, A., Heidrich, L., Siobal, M., Isensee, J., Prolingheuer, J., Janocha, E., Valdor, M., Hagendorf, S., Bahrenberg, G., Opitz, T., Segschneider, M., Haupt, S., Nitzsche, A., Brüstle, O., & Hucho, T. (2023). Emergence of nociceptive functionality and opioid signaling in human induced pluripotent stem cell–derived sensory neurons. PAIN. https://journals.lww.com/pain/Fulltext/9900/Emergence_of_nociceptive_functionality_and_opioid.249.aspx

Schwartzentruber, J., Foskolou, S., Kilpinen, H., Rodrigues, J., Alasoo, K., Knights, A. J., Patel, M., Goncalves, A., Ferreira, R., Benn, C. L., Wilbrey, A., Bictash, M., Impey, E., Cao, L., Lainez, S., Loucif, A. J., Whiting, P. J., Gutteridge, A., Gaffney, D. J., & HIPSCI Consortium, D. J. (2018). Molecular and functional variation in iPSC-derived sensory neurons. 50(1), 54–61. 10.1038/s41588-017-0005-8

Shiers, S. I., Mazhar, K., Wangzhou, A., Haberberger, R., Lesnak, J. B., Sankaranarayanan, I., Tavares-Ferreira, D., Cervantes, A., Funk, G., Horton, P., Vines, E., Dussor, G., & Price, T. J. (2024). Nageotte nodules in human DRG reveal neurodegeneration in painful diabetic neuropathy. BioRxiv, 2024.08.22.609215. 10.1101/2024.08.22.609215

Stuart, T., Butler, A., Hoffman, P., Hafemeister, C., Papalexi, E., Mauck, W. M., Hao, Y., Stoeckius, M., Smibert, P., & Satija, R. (2019). Comprehensive Integration of Single-Cell Data. Cell, 177(7), 1888–1902.e21. 10.1016/J.CELL.2019.05.031

Sun, Y., Dykes, I. M., Liang, X., Eng, S. R., Evans, S. M., & Turner, E. E. (2008). A central role for Islet1 in sensory neuron development linking sensory and spinal gene regulatory programs. Nature Neuroscience, 11(11). http://www.ncbi.nlm.nih.gov/pubmed/18849985

Tavares-Ferreira, D., Shiers, S., Ray, P. R., Wangzhou, A., Jeevakumar, V., Sankaranarayanan, I., Cervantes, A. M., Reese, J. C., Chamessian, A., Copits, B. A., Dougherty, P. M., Gereau, R. W., Burton, M. D., Dussor, G., & Price, T. J. (2022). Spatial transcriptomics of dorsal root ganglia identifies molecular signatures of human nociceptors. Science Translational Medicine, 14(632), eabj8186. 10.1126/SCITRANSLMED.ABJ8186/SUPPL_FILE/SCITRANSLMED.ABJ8186_FILES_S1_TO_S31.ZIP

Thomas, P. D., Ebert, D., Muruganujan, A., Mushayahama, T., Albou, L. P., & Mi, H. (2022). PANTHER: Making genome-scale phylogenetics accessible to all. Protein Science, 31(1), 8–22. 10.1002/PRO.4218

Usoskin, D., Furlan, A., Islam, S., Abdo, H., Lönnerberg, P., Lou, D., Hjerling-Leffler, J., Haeggström, J., Kharchenko, O., Kharchenko, P. V., Linnarsson, S., & Ernfors, P. (2015). Unbiased classification of sensory neuron types by large-scale single-cell RNA sequencing. Nature Neuroscience, 18(1), 145–153. 10.1038/nn.3881

Vermeiren, S., Bellefroid, E. J., & Desiderio, S. (2020). Vertebrate Sensory Ganglia: Common and Divergent Features of the Transcriptional Programs Generating Their Functional Specialization. Frontiers in Cell and Developmental Biology, 8, 587699. 10.3389/FCELL.2020.587699/BIBTEX

Waddell, P. J., S.N., L., & McCarthy, P. W. (1989). Conduction velocity changes along the processes of rat primary sensory neurons. Neuroscience, 30(3), 577–584. 10.1016/0306-4522(89)90152-8

Wickham, H. (2016). ggplot2 - Elegant Graphics for Data Analysis. Springer International Publishing. 10.1007/978-3-319-24277-4

Xiao, D., Deng, Q., Guo, Y., Huang, X., Zou, M., Zhong, J., Rao, P., Xu, Z., Liu, Y., Hu, Y., Shen, Y., Jin, K., & Xiang, M. (2020). Generation of self-organized sensory ganglion organoids and retinal ganglion cells from fibroblasts. Science Advances, 6(22), eaaz5858. 10.1126/sciadv.aaz5858

Yu, G., Wang, L. G., Han, Y., & He, Q. Y. (2012). ClusterProfiler: An R package for comparing biological themes among gene clusters. OMICS A Journal of Integrative Biology, 16(5), 284–287. 10.1089/OMI.2011.0118/ASSET/IMAGES/LARGE/FIGURE1.JPEG

Zhang, C.-J., Ji, M.-J., Zhou, X.-L., Li, S., Xu, P.-F., Wu, H., Chen, Q., Zhao, J., Chen, X.-Z., Cox, J. J., & Zhou, X.-L. (2026). Nav1.5 in the dorsal root ganglion plays a crucial role in mechanical hypersensitivity. Brain, 149(7), 2516–2532. 10.1093/BRAIN/AWAF438

Zhang, Y., Pak, C. H., Han, Y., Ahlenius, H., Zhang, Z., Chanda, S., Marro, S., Patzke, C., Acuna, C., Covy, J., Xu, W., Yang, N., Danko, T., Chen, L., Wernig, M., & Südhof, T. C. (2013). Rapid Single-Step Induction of Functional Neurons from Human Pluripotent Stem Cells. Neuron, 78(5), 785. 10.1016/J.NEURON.2013.05.029

Zhao, Y., Wang, X., & Wang, K. (2023). Transcription factor-mediated programming of stem cell fate. Trends in Cell Biology, 33(8), 621–624. 10.1016/j.tcb.2023.05.004

Zheng, Y., Liu, P., Bai, L., Trimmer, J. S., Bean, B. P., & Ginty, D. D. (2019). Deep Sequencing of Somatosensory Neurons Reveals Molecular Determinants of Intrinsic Physiological Properties. Neuron, 103(4), 598–616.e7. 10.1016/j.neuron.2019.05.039

