## Supplementary Figures for "In vitro programming and pseudounipolarization of human iPSC-derived sensory neurons"

### Suppl. Figure 1

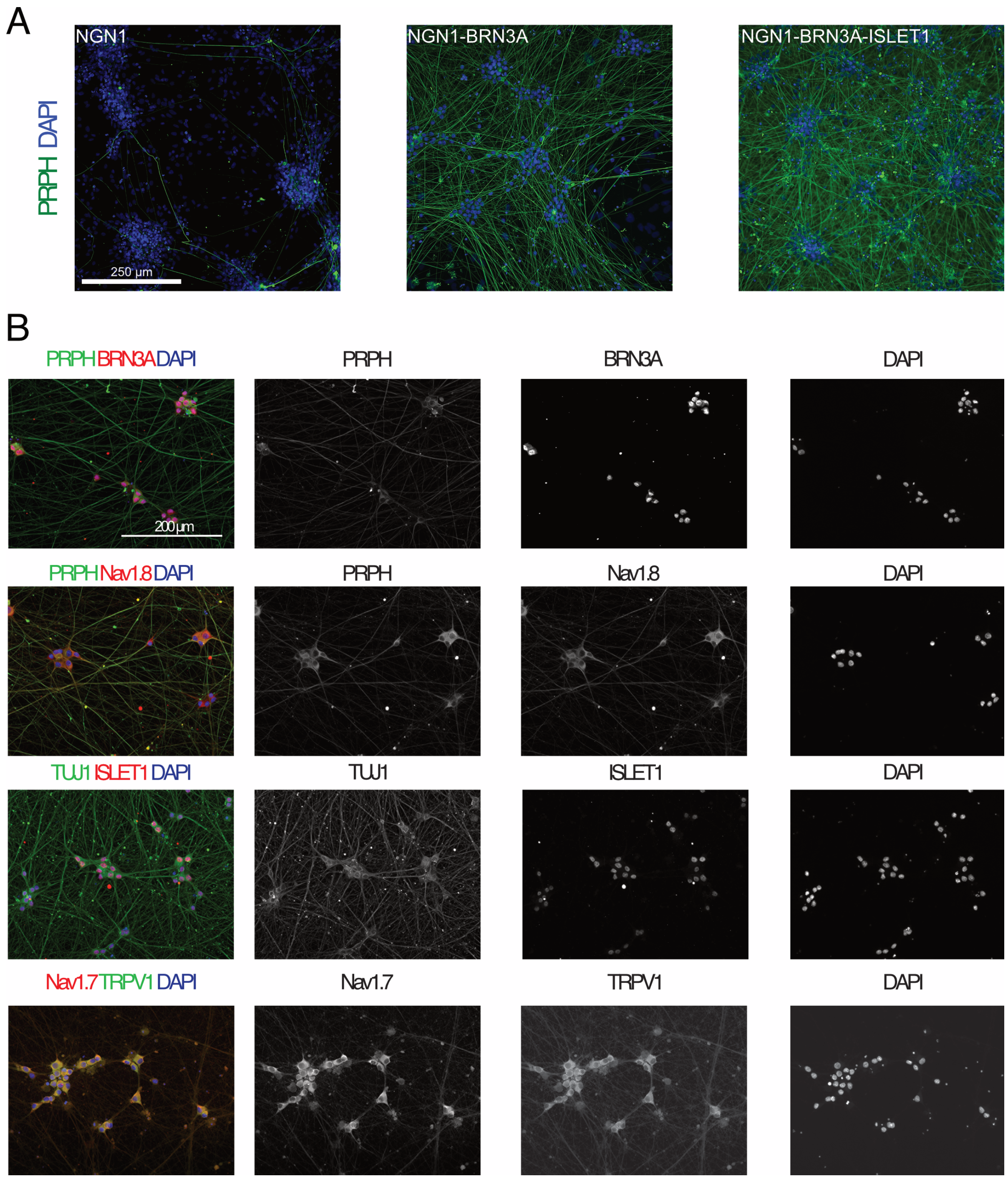


**Supplementary Figure 1. Overexpression of NBI efficiently generates PRPH+ neurons.**
**(A)** Immunohistochemical staining of differentiated cultures after overexpression of NGN1, NB or NBI for one week for the sensory neuron marker peripherin. **(B)** Representative images of sensory neuron marker expression in iNBI-SN cultures at DIV 63 as shown in Figure 1, supplemented by single channel images depicted in gray scale.

### Suppl. Figure 2

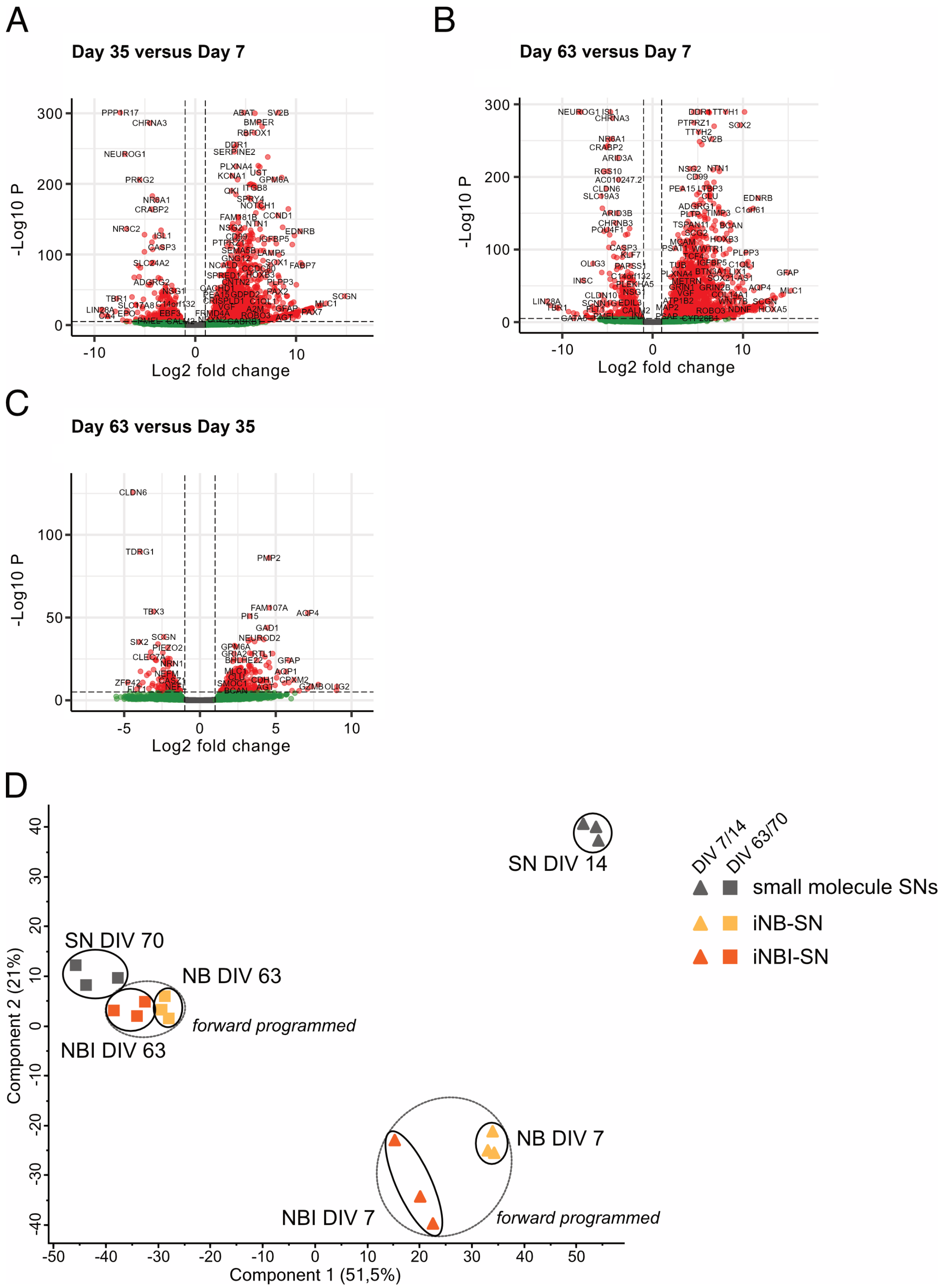


**Supplementary Figure 2. Transcriptome and proteome analysis of iPSC-derived sensory neurons.**
**(A-C)** Volcano plots illustrating differential gene expression between time points DIV7 and DIV35 (A), DIV7 and DIV63 (B) and DIV35 and DIV63 (C). **(D)** Principal component analysis (PCA), illustrating the clustering of proteome data of iPSC-derived sensory neurons generated via a classical small

molecule protocol (dual SMAD inhibition) or forward programming (iNB-SN, iNBI-SN) across two principal components, accounting for 51.5 % (PC1) and 21.0 % (PC2) of total variance. Protein expression was analyzed after initial fate induction i.e., DIV 14 for the small molecule protocol and DIV 7 for the forward programming protocol and after prolonged maturation in the presence of neurotrophic factors NGF, BDNF, GDNF and NT-3 at DIV 70 and 63, respectively. Populations were encircled for better visualization.

**Suppl. Figure 3**

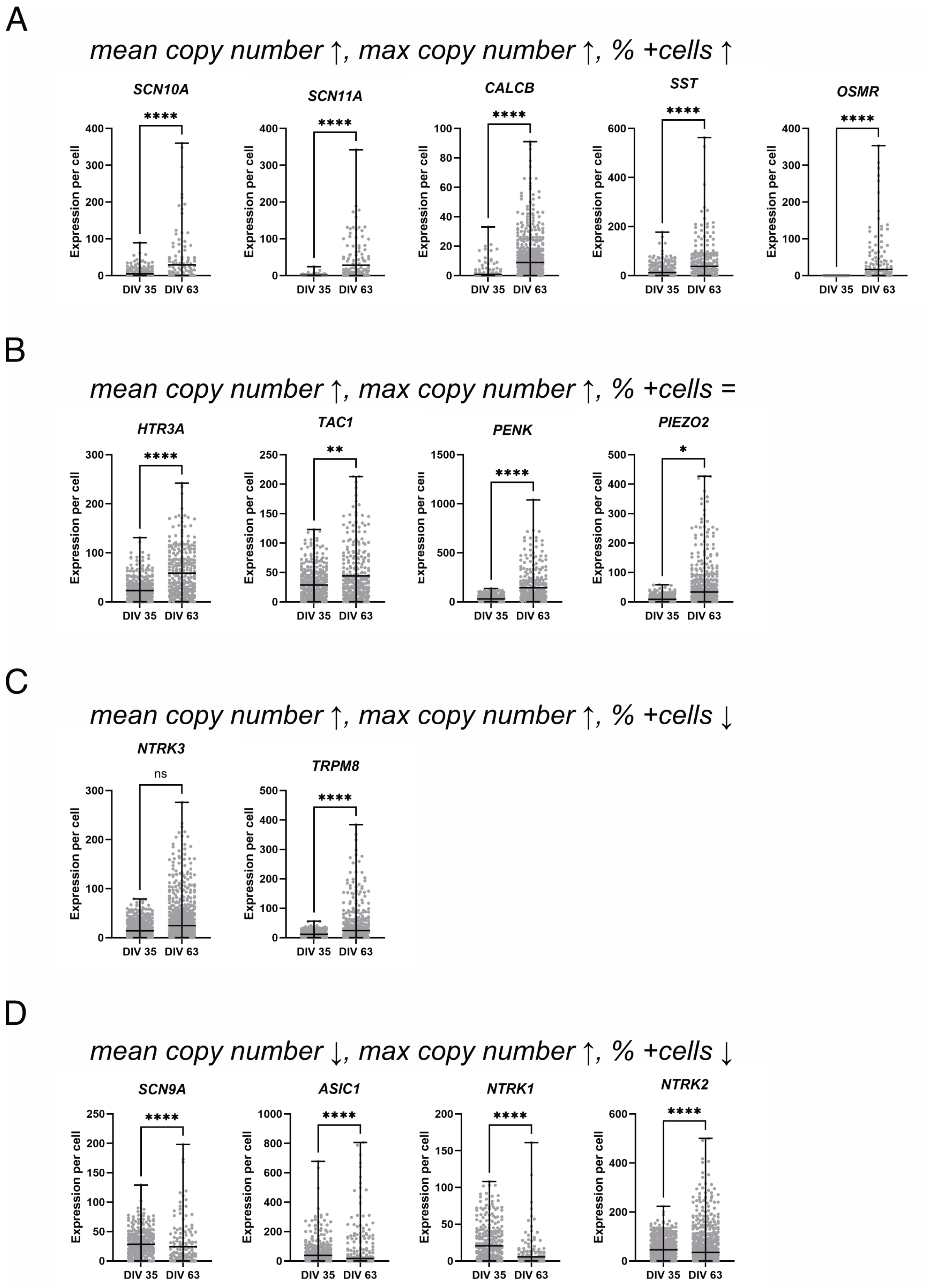


**Supplementary Figure 3.** RNAscope quantification of gene expression per cell at DIV 35 and DIV 63. **(A)** Targets (*SCN10A, SCN11A, CALCB, SST, OSMR*) that showed an increased mean copy number per cell and increased percentage of expressing cells. **(B)** Targets (*HTR3A, TAC1, PENK, PIEZO2*) that showed an increased mean copy number per cell, but no change in percentage of expressing cells. **(C)** Targets (*NTRK3, TRPM8*) that showed an increased mean copy number per cell and a reduction in percentage of expressing cells. **(D)** Targets (*SCN9A, ASIC1, NTRK1, NTRK2*) that showed a decreased mean copy number per cell and a reduction in percentage of expressing cells. Values are presented as mean with range and single data points indicated by circles. Mann-Whitney test.

**Suppl. Figure 4**

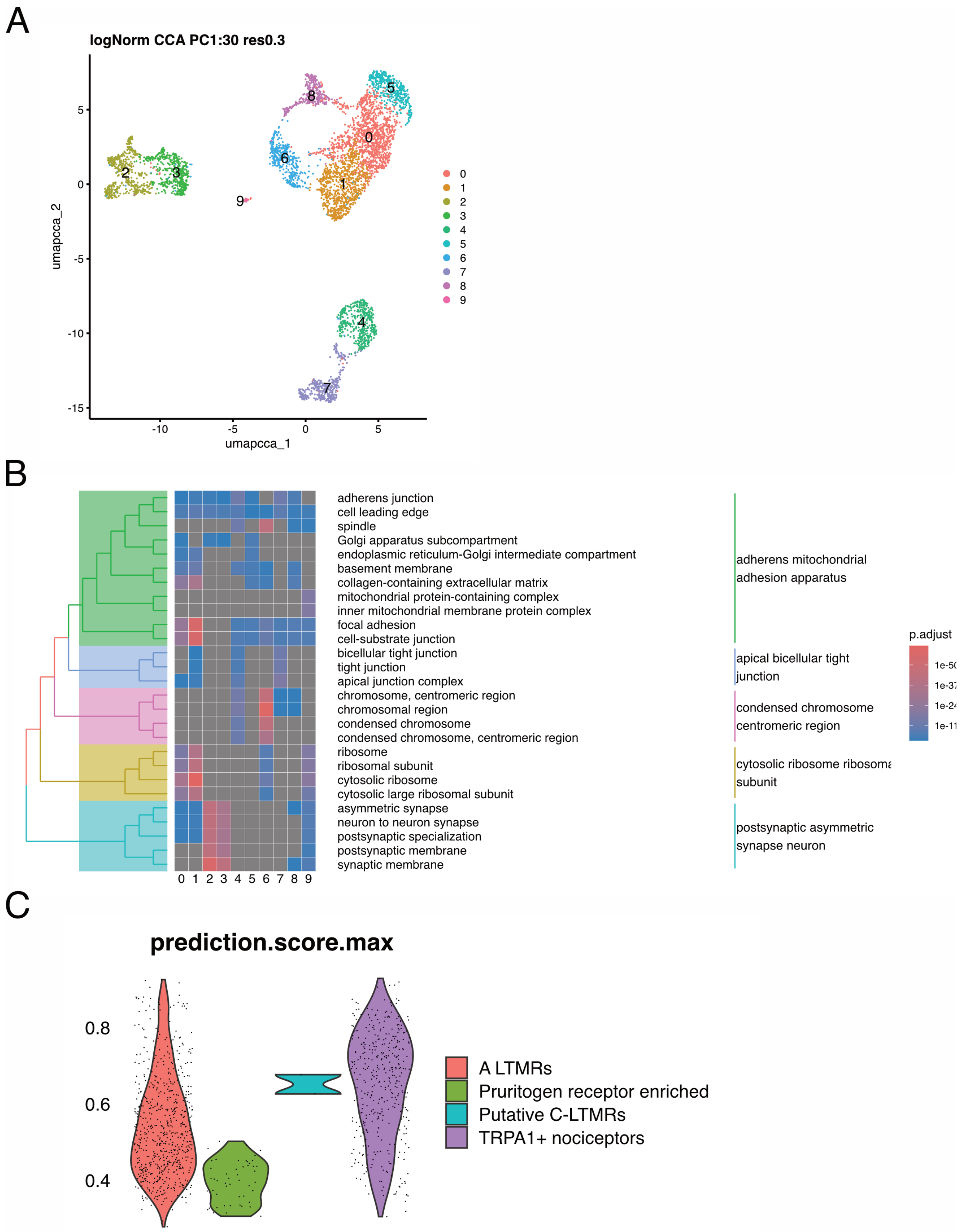


**Supplementary Figure 4. Single nucleus RNA sequencing analysis.**
**(A)** UMAP plot with clusters identified from all cells in iNBI-SN cultures at DIV 63 derived from two healthy donors. **(B)** Gene ontology enrichment analysis of identified clusters. **(C)** Violin plots display the distribution of prediction scores for each transferred label.

**Suppl. Figure 5**

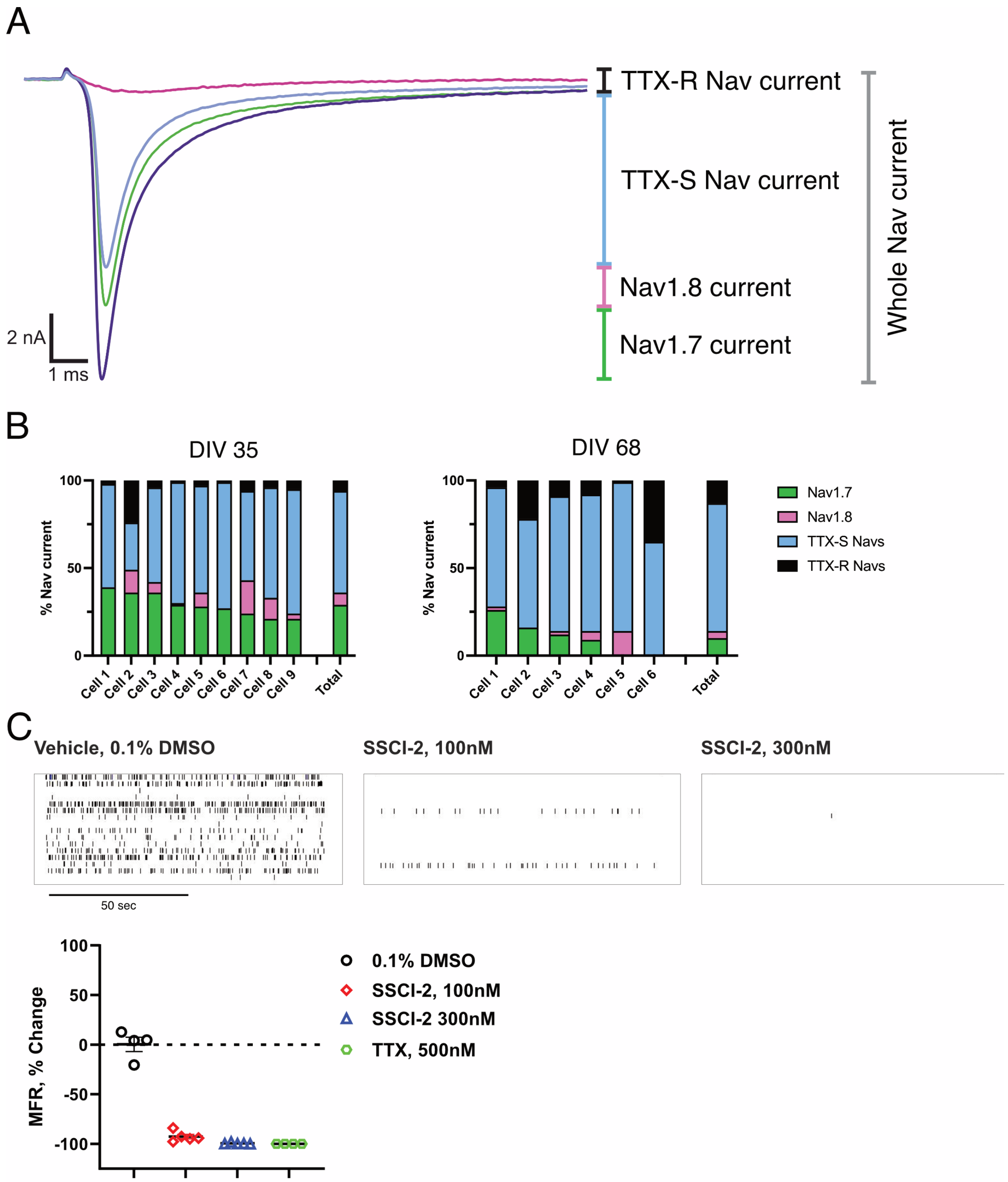


**Supplementary Figure 5. Sodium current analysis of iNBI-SNs.**
**(A)** Exemplary sodium current trace of iNBI-SN at DIV 35, indicating the contribution of Nav1.7 and Nav1.8, as well as the remaining TTX-S and TTX-R currents to the total sodium current. Sequential application of 100 nM SSCI-2, 100 nM SSCI-2 + 3 µM VX-150 and 100 nM SSCI-2 + 3 µM VX-150 + 500 nM TTX were employed to determine Nav1.7, Nav1.8 and the remaining TTX-R current, respectively. **(B)** Quantification of the proportion of Nav1.7, Nav1.8, TTX-S and TTX-R to the total sodium current at DIV 35 and 68. **(C)** Spike firing raster plot for vehicle, 100 nM SSCI-2 and 300 nM SSCI-2 treated iNBI-SN cultures. Raster plot indicates reduction of spontaneous firing both at 100 nM and 300 nM as compared to vehicle control. Quantification of the change in mean firing rate upon treatment compared to baseline at DIV 64. N=4-5 wells. Data are presented as mean ± SEM.

**Suppl. Figure 6**

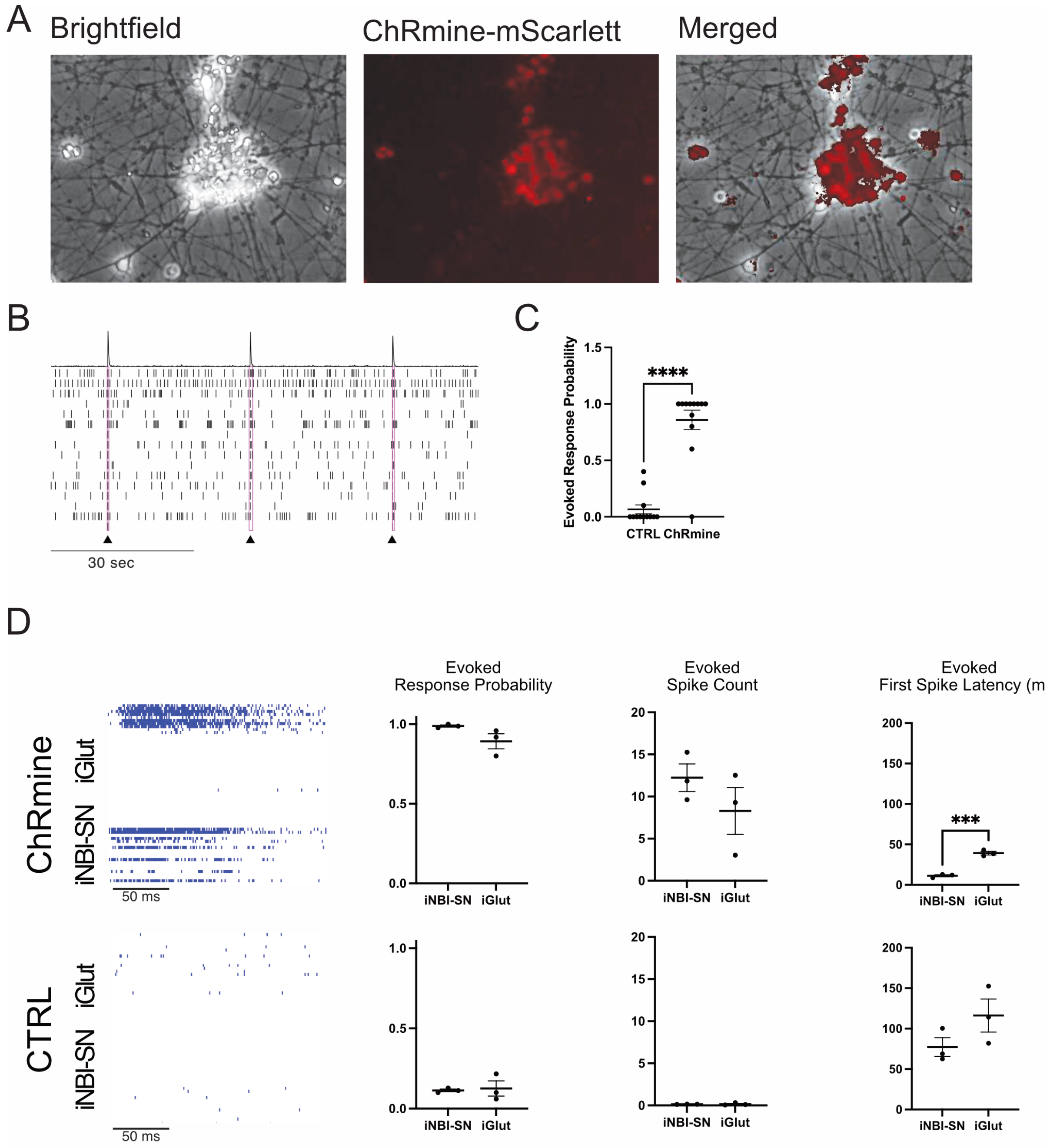


**Supplementary Figure 6. Transmission of optogenetically evoked signals from iNBI-SN to iGlut neurons.**

**(A)** Representative brightfield and fluorescence microscopy images of iNBI-SNs at DIV 7 after transduction with AAVs introducing the optogenetic tool ChRmine, including a red fluorescent mScarlett tag. Expression of ChRmine was apparent in the majority of iNBI-SNs and was well tolerated. **(B)** Spike raster plot of baseline and optically evoked activity. Black triangle indicates optical stimulation of iNBI-SN culture equipped with ChRmine. **(C)** Evoked response probability of control iNBI-SNs and iNBI-SNs that express ChRmine. ChRmine expressing iNBI-SNs showed a clear activation and an evoked response probability of more than 85%. N=12 wells. Data are presented as mean ± SEM. Mann-Whitney test. **(D)** To test the functional transmission of an electrical signal from iNBI-SNs towards CNS neurons, iGlut neurons were paired with iNBI-SNs equipped with the optogenetic activator ChRmine, or control iNBI-SNs. Raster plot of evoked spiking behavior upon optical stimulation in ChRmine and control iNBI-SN and iGlut co-cultures. Quantification of the evoked response probability, evoked spike count and evoked first spike latency upon optical stimulation. N=3 wells with 32 electrodes / neuron type / well. Values are presented as mean ± SEM. Unpaired t-test.

**Table S1: qRT-PCR primer**

| Gene | Forward primer | Reverse primer |
|---|---|---|
| GAPDH | GGCCTCCAAGGAGTAAGACC | AGGGGTCTACATGGCAACTG |
| Ki67 | AGGGAAAGGAGAAGCAGGAA | GACACACACATTGTCCTCAGC |
| NTRK1 | CGCTCTCGGTGGTGAACT | CAGGGACTGGTGGTCAAGAT |
| NTRK2 | TGTTCAGCACATCAAGCGACA | CAAAGGCTCCTTCGCCTAGC |
| NTRK3 | GCGTTTCAAAGAAGCAGCGAT | CGGCAATTGATCTCAGTCTTGC |
| OCT4/ POU5F1 | GTGGAGGAAGCTGACAACAA | ATTCTCCAGGTTGCCTCTCA |
| PRPH | TGATCAAGACCATTGAGACCCG | TCAGTAACTGTGGGCAGAAGAC |
| SCN9A | GTCCAAGAAGCCACAAAAGC | CTTTGACCCTCCTTTTCTACCA |
| SCN11A | GGCAAGAGGTTTCATTCTGG | CAGGGGCAATAGTTTGATGG |
| TRPV1 (Namer et al., 2019) | GCACAGGAGAGCAAGAACATC | GTCCAGTTCACCTCGTCCAC |

**Table S2 Antibodies**

| Antibody | Company | Catalogue | Dilution |
|---|---|---|---|
| rabbit anti-βIII-tubulin | Synaptic Systems | 302302 | 1:1000 |
| mouse anti-human nuclei | Merck | MAB4383 | 1:100 |
| Mouse anti PRPH | Santa Cruz Biotechnology | Sc-37709-3 | 1:200 |
| S100b | abcam | ab52642 | 1:200 |
| BRN3A | Merck | AB5945 | 1:200 |
| ISLET1 | Abcam | ab86501 | 1:100 |
| rabbit anti-βIII-tubulin | BioLegend | 802001 | 1:1000 |
| Nav1.7 | abcam | ab85015 | 1:100 |
| Nav1.8 | Alomone labs | ASC-016 | 1:100 |
| Chicken anti MAP2 | Novus bio | #NB300-213 | 1:1000 |
| Goat anti chicken IgG-Alexa Fluor 647 | Thermo Fisher | A-21449 | 1:500 |
| donkey anti rabbit IgG-Alexa Fluor 488 | Thermo Fisher | A-21206 | 1:500 |
| donkey anti mouse IgG-Alexa Fluor 594 | Thermo Fisher | A-21203 | 1:500 |
| Alexa Fluor® 594 AffiniPure Donkey Anti-Rabbit | Jackson ImmunoResearch Labs | 711-585-152 | 1:500 |
| Alexa Fluor® 647 AffiniPure Donkey Anti-Mouse | Jackson ImmunoResearch Labs | 715-605-150 | 1:500 |

**Table S3: RNAscope probes**

| mRNA | Channel | Catalogue number | Fluorophore (1/1000) dilution |
|---|---|---|---|
| *SCN9A* | C1 | 562251 | TSA 520 |
| *SCN10A* | C2 | 406291-C2 | TSA 570 |
| *SCN11A* | C3 | 404791-C3 | TSA 650 |

| | | | |
|---|---|---|---|
| *ASIC1* | C1 | 576251 | TSA 520 |
| *NTRK2* | C2 | 402621-C2 | TSA 570 |
| *NTRK3* | C3 | 406341-C3 | TSA 650 |

| | | | |
|---|---|---|---|
| *HTR3A* | C1 | 310681 | TSA 520 |
| *TAC1* | C2 | 310711-C2 | TSA 570 |
| *NTRK1* | C3 | 402631-C3 | TSA 650 |

| | | | |
|---|---|---|---|
| *CALCB* | C1 | 849281 | TSA 520 |
| *PIEZO2* | C2 | 449951-C2 | TSA 570 |
| *TRPM8* | C3 | 543121-C3 | TSA 650 |

| | | | |
|---|---|---|---|
| *PENK* | C1 | 548301 | TSA 520 |
| *OSMR* | C2 | 537121-C2 | TSA 570 |
| *SST* | C3 | 310591-C3 | TSA 650 |

| | | | |
|---|---|---|---|
| 3-plex negative Control | C1 | 320871 | TSA 520 |
| | C2 | | TSA 570 |
| | C3 | | TSA 650 |

| | | | |
|---|---|---|---|
| 3-plex positive Control | C1 | 320861 | TSA 520 |
| | C2 | | TSA 570 |
| | C3 | | TSA 650 |
